# Molecular basis for antiviral activity of pediatric neutralizing antibodies targeting SARS-CoV-2 Spike receptor binding domain

**DOI:** 10.1101/2022.07.27.501708

**Authors:** Yaozong Chen, Jérémie Prévost, Irfan Ullah, Hugo Romero, Veronique Lisi, William D Tolbert, Jonathan R. Grover, Shilei Ding, Shang Yu Gong, Guillaume Beaudoin-Bussières, Romain Gasser, Mehdi Benlarbi, Dani Vézina, Sai Priya Anand, Debashree Chatterjee, Guillaume Goyette, Michael W. Grunst, Ziwei Yang, Yuxia Bo, Fei Zhou, Kathie Béland, Xiaoyun Bai, Allison R. Zeher, Rick K. Huang, Dung N. Nguyen, Rebekah Sherburn, Di Wu, Grzegorz Piszczek, Bastien Paré, Doreen Matthies, Di Xia, Jonathan Richard, Priti Kumar, Walther Mothes, Marceline Côté, Pradeep D. Uchil, Vincent-Philippe Lavallée, Martin A. Smith, Marzena Pazgier, Elie Haddad, Andrés Finzi

## Abstract

Neutralizing antibodies (NAbs) hold great promise for clinical interventions against SARS-CoV- 2 variants of concern (VOCs). Understanding NAb epitope-dependent antiviral mechanisms is crucial for developing vaccines and therapeutics against VOCs. Here we characterized two potent NAbs, EH3 and EH8, isolated from an unvaccinated pediatric patient with exceptional plasma neutralization activity. EH3 and EH8 cross-neutralize the early VOCs and mediate strong Fc-dependent effector activity *in vitro*. Structural analyses of EH3 and EH8 in complex with the receptor-binding domain (RBD) revealed the molecular determinants of the epitope-driven protection and VOC-evasion. While EH3 represents the prevalent IGHV3-53 NAb whose epitope substantially overlaps with the ACE2 binding site, EH8 recognizes a narrow epitope exposed in both RBD-up and RBD-down conformations. When tested *in vivo*, a single-dose prophylactic administration of EH3 fully protected stringent K18-hACE2 mice from lethal challenge with Delta VOC. Our study demonstrates that protective NAbs responses converge in pediatric and adult SARS-CoV-2 patients.

## INTRODUCTION

The Coronavirus Disease 2019 (COVID-19) pandemic, caused by the betacoronavirus SARS-CoV-2, has resulted in more than 577 million infections and over 6.4 million deaths worldwide as of July 2022. The rapid development and implementation of vaccines has proven to be a remarkable success in controlling the global pandemic. However, since the end of 2020, a number of SARS-CoV-2 mutants that can escape vaccine or infection induced immune responses have emerged worldwide, designated variants of concern (VOCs). These now include SARS-CoV-2 strains Alpha (B.1.1.7) ^1^, Beta (B.1.351) ^2^, Gamma (P.1) ^3^, Delta (B.1.617.2) ^4^ and most recently Omicron (B.1.1.529) BA.1-BA.5 ^5^, which currently pose a threat to global recovery. Novel prophylactic and therapeutic interventions are needed to combat future outbreaks by these and other SARS-CoV-2 variants.

Children with SARS-CoV-2 infection may experience different clinical symptoms from adults, including a greater chance of being asymptomatic, more persistent low-level infections and higher ratio of gastrointestinal upset over respiratory illness ^6^. These differences could be a consequence of lower anti-nucleocapsid IgG responses or reduced serum neutralizing activity in children ^7^. How these differences compare to other age groups and whether they affect the magnitude or specificity of antibody responses to other SARS-CoV-2 antigens is still a matter of debate. One of the more commonly used heavy chain variable genes for RBD-targeting antibodies in SARS-CoV-2 seropositive adult subjects is IGHV3-53 ^8, 9^. IGHV3-53 encoded antibodies specific for SARS-CoV-2 show potent neutralization and have been referred to as “public antibodies” ^9^. They are characterized by relatively short heavy chain Complementary Determining Region 3s (CDRH3s), a low level of somatic mutation, as well as a similar epitope footprint and angle of approach to the RBD that directly overlaps the ACE2 footprint ^8–10^. This disproportionate convergence of certain variable genes in response to infection has also been reported in other pathogens such as dengue virus ^11^, HIV-1 ^12^, malaria ^13^ and influenza ^14^. Interestingly, a single RBD mutation K417N/T present in B.1.351, P.1 and Omicron variants has been reported to confer resistance to these IGHV3-53 encoded antibodies ^9^, consistent with the hypothesis that immune selection by antibodies prevalent in the population is a vital attribute driving viral evolution ^15–17^.

Existing evidence supports the notion that antibody-mediated protection against viral infections relies on several functional components. Direct neutralizing activities constitute the first line of defense mostly by preventing viral entry into target cells. The second line of protection is more complex and is composed of variable Fc-mediated effector functions that eliminate virus and virally infected cells including antibody-dependent cellular cytotoxicity (ADCC) mediated mostly by natural killer (NK) cells–and antibody-dependent cellular phagocytosis (ADCP) mediated by neutrophils, monocytes macrophages and dendritic cells. Several recent studies using animal models clearly indicate that *in vivo* efficacies of anti-SARS-CoV-2 antibodies rely on a combination of neutralization and Fc-effector activities ^18–21^, with the latter playing an indispensable role in disease mitigation. Like neutralization, precise epitope targeting and binding affinity were found to be major determinants contributing to the effectiveness of Fc-effector mechanisms of SARS-CoV-2 Spike-targeting monoclonal Antibodies (mAbs) ^22^.

While most anti-SARS-CoV-2 antibodies characterized to date include antibodies isolated from adult patients, less is known about the humoral response induced by infection in the pediatric population. Here, we isolated two potent neutralizing antibodies, EH3 and EH8, from an unvaccinated pediatric patient in the early onset of the COVID-19 pandemic. EH3 is encoded by IGHV3-53/IGKV3-20, representing the most prevalent class of anti-RBD antibodies isolated from SARS-CoV-2 adult patients, while EH8 is encoded by the rarely used IGHV1-18/IGLV2-23 variable genes. Both antibodies bind the SARS-CoV-2 RBD and its variants with low nanomolar affinity and effectively neutralize pseudoviral particles bearing VOCs Spikes (except for Omicron). Structural analysis provided an atomic understanding of antibody-RBD interactions: EH3 resembles a typical IGHV3-53 encoded antibody whose epitope highly overlaps with the ACE2 binding site, while EH8 binds to the narrow RBD ridge region that is fully exposed in the RBD-down Spike. Moreover, we further assessed EH3 for its prophylactic potential in a stringent K18-hACE2 mouse model challenged by the Delta variant, which validated its robust *in-vivo* efficacy against lethal SARS-CoV-2 infection. These data indicate that protective RBD-specific IGHV3-53 encoded antibodies can also be found in pediatric patients. Furthermore, although induced in response to infection with the early circulating SARS-CoV-2 (Wuhan-Hu-1 strain, WT) strain, this antibody class preserves its antiviral activity against all tested VOCs except Omicron.

## RESULTS

### Isolation of RBD-specific NAbs from an unvaccinated pediatric COVID-19 patient

In the early onset of COVID-19 pandemic (April 2020), plasma from a 6-year-old patient with multisystemic inflammatory syndrome in children (MIS-C) was analysed for the presence of neutralizing antibodies able to block viral entry using a well-established neutralization assay based on pseudoviral particles bearing the SARS-CoV-2 Spike^23^. High neutralization titers were found in plasma from this pediatric patient (**Figure 1A**), which were higher than previously characterized adult elite neutralizers (S002, S006)^24^. Since the most dominant neutralizing antibody responses target RBD, this plasma sample was tested for the presence of anti-RBD antibodies using a previously described ELISA assay^23^. In line with its neutralization activity, we found that this patient had high levels of anti-RBD IgG as compared to adult neutralizers (**Figure 1B**). Based on these results, we sought to characterize the patient’s anti-RBD B cell repertoire in detail. To do so, we isolated peripheral blood mononuclear cells (PBMCs) from a limited blood sample (∼5mL) and performed cell-surface staining to identify RBD-specific B cell clones (**Figure 1C**). A total of nine RBD^+^ CD19^+^ IgG^+^ IgD^-^ CD3^-^ CD14^-^ B cells were sorted (**Figure S1**), and five clones were successfully sequenced, assembled, cloned and expressed in 293F cells. The neutralization potencies of the purified anti-RBD IgGs were initially tested on pseudoviruses expressing the autologous SARS-CoV-2 S WT (Wuhan-Hu-1) strain. EH3 and EH8 were the top neutralizers of the group with half maximal inhibitory concentrations (IC50) in the range of ∼4-13 ng/mL (**Figure 1D**). Lineage analysis for heavy and light chain genes indicated that EH3 was encoded by the IGHV3-53 heavy chain variable gene, paired with an IGKV3-20 encoded light chain gene, which represents a prevalent class of RBD-specific antibodies found in many SARS-CoV-2 patients^8, 9^ (**Figure S4A**). EH8, on the other hand, is composed of two rarely used germline variable genes in SARS-CoV-2 infection, IGHV1-18 and IGLV2-23 (**Figure S4B**).

**Figure 1.**
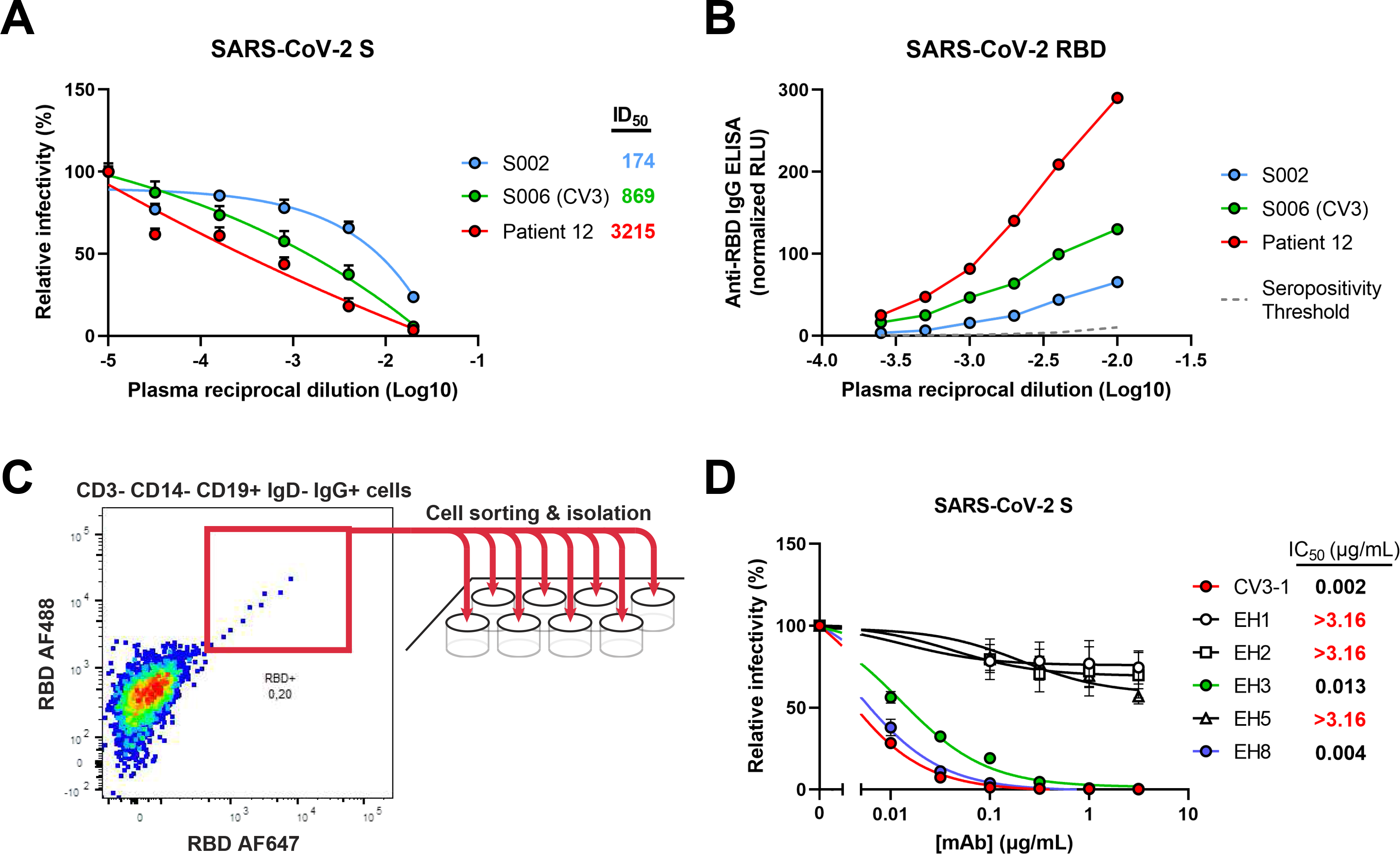
Isolation of RBD-specific mAbs from a pediatric patient. (**A**) Pseudoviruses encoding the luciferase gene (Luc+) and bearing SARS-CoV-2 full-length S (Wuhan-Hu-1 strain) were used to infect 293T-hACE2 cells in presence of increasing dilutions of indicated COVID-19+ plasma samples at 37 °C for 1 h prior infection of 293T-hACE2 cells. Fitted curves and half maximal inhibitory dilution (ID50) values were determined using a normalized nonlinear regression. (**B**) Indirect ELISA was performed using recombinant SARS-CoV-2 RBD protein and incubation with COVID-19+ plasma samples from two adult donors (S002, S006 [CV3]) and the pediatric patient of interest (Patient 12). Anti-RBD antibody binding was detected using horseradish peroxidase (HRP)-conjugated anti-human IgG. Relative light unit (RLU) values obtained with BSA (negative control) were subtracted and further normalized to the signal obtained with the anti-RBD mAb CR3022 present in each plate. Seropositivity thresholds were calculated using ten pre-pandemic COVID-19-negative plasma samples. (**C**) Cryopreserved PBMCs obtained from Patient 12 were stained for the expression of cell-surface markers (CD3, CD14, CD19, IgD, IgG) and probed with fluorescently-labelled SARS-CoV-2 RBD proteins. RBD-specific B cells (CD3- CD14- CD19+ IgD- IgG+ RBD-AF488+ RBD-AF647+) were individually sorted into a 96-well plate, followed by targeted B-cell receptor sequencing. (**D**) Pseudoviruses Luc+ bearing SARS-CoV-2 full-length S (Wuhan-Hu-1 strain) were used to infect 293T-hACE2 cells in presence of increasing concentrations of indicated mAbs isolated from Patient 12 at 37°C for 1 h prior infection of 293T-hACE2 cells. Fitted curves and half maximal inhibitory antibody concentration (IC50) values were determined using a normalized nonlinear regression. Error bars indicate means ± SEM.

### EH3 and EH8 effectively bind and neutralize most SARS-CoV-2 VOCs

First, we characterized the neutralizing activities and Fc-effector functions of these newly identified pediatric NAbs using the highly potent anti-RBD NAb CV3-1 as a control (**Figure 1D**); CV3-1 was previously isolated from the S006 donor (also known as CV3) and thoroughly characterized for its antiviral activities both *in vitro* and *in vivo* ^21, 25^. We examined the cross-reactivity of EH3 and EH8 to recognize and neutralize the emerging SARS-CoV-2 VOCs and variants of interest (VOIs), including B.1.1.7 (Alpha), B.1.351 (Beta), P.1 (Gamma), B.1.429 (Epsilon), B.1.526 (Iota), B.1.617.1 (Kappa), B.1.617.2 (Delta) and B.1.1.529 (Omicron BA.1 and BA.2). EH3 efficiently bound to cells and neutralized pseudoviral particles bearing S glycoproteins from all VOCs and VOIs with IC50 ranging from 13-34 ng/mL, except for the B.1.1.529 lineages (BA.1 and BA.2) (**Figures 2A, C**). On the other hand, EH8 binding and neutralizing breadth was limited to early VOCs and VOIs, while its recognition and neutralization of the Delta (B.1.617.2) and Omicron variants (B.1.1.529.1 and B.1.1.529.2) were substantial reduced (**Figures 2B, D**). Although their neutralization potency against pseudoviruses was similar (**Figures 1D, 2C-D**), EH8 was 3 times more potent in inhibiting S-driven cell-to-cell fusion than EH3 (**Figure 2E**). To evaluate Fc-mediated effector functions, we used an assay that quantifies the antibody-dependent cellular cytotoxicity (ADCC) activity against SWT-expressing cells. While we did not observe obvious differences in antibody binding levels or cooperativity (**Figure 2F**), EH8 mediated the elimination of S-expressing cells more efficiently than EH3 at higher antibody concentrations (**Figure 2G**).

**Figure 2.**
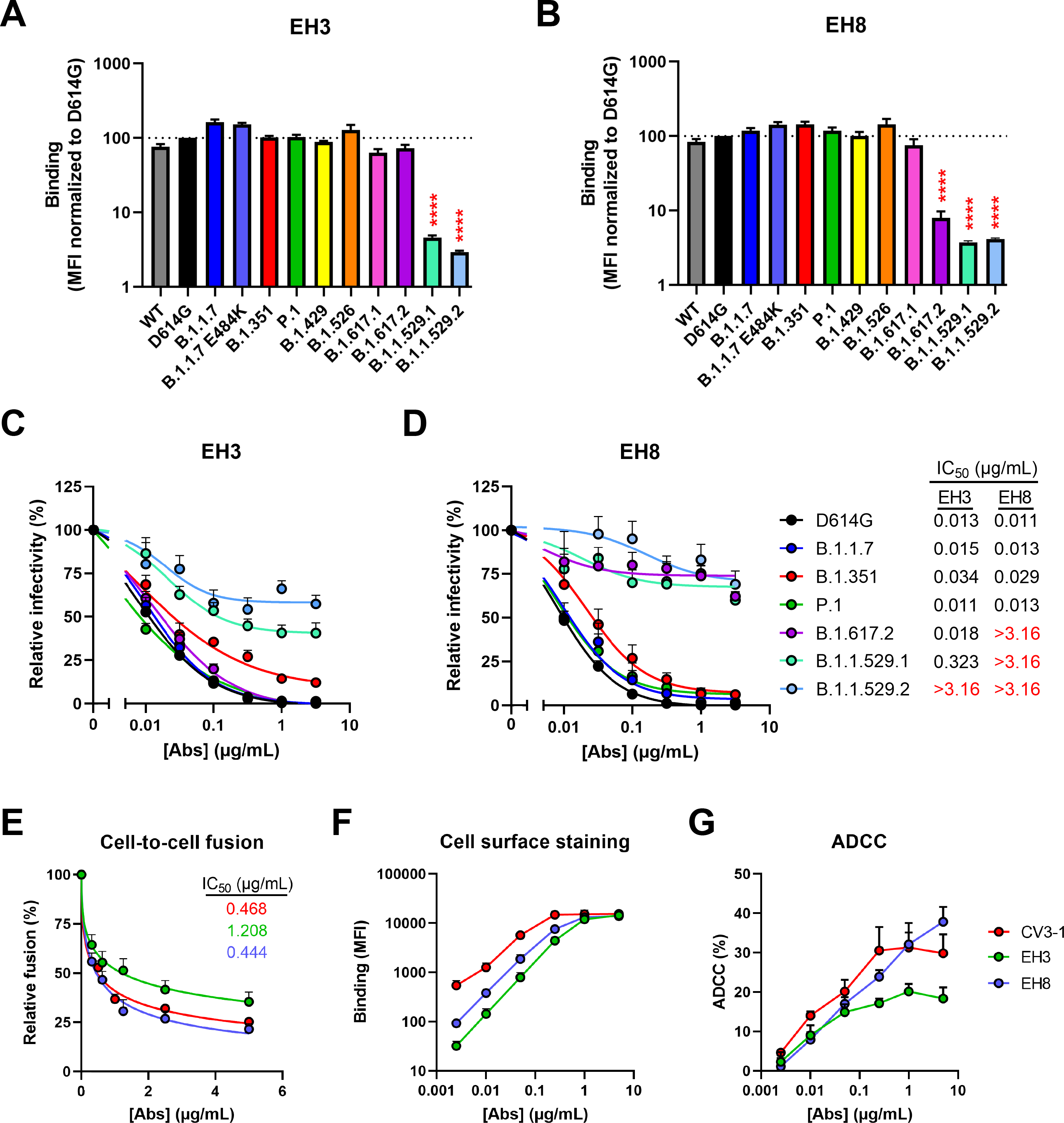
Characterization of potent pediatric RBD-specific neutralizing mAbs. (**A-B**) Cell-surface staining of 293T cells expressing full-length S from indicated variants using EH3 (**A**) and EH8 (**B**) mAbs. The graphs show the median fluorescence intensities (MFI). Dashed lines indicate the reference value obtained with S D614G. Statistical significance was tested using mixed-effects ANOVA with a Dunnett post-test (∗∗∗∗, p <0.0001). (**C-D**) Pseudoviruses encoding the luciferase gene (Luc+) and bearing SARS-CoV-2 full-length S from indicated variants were used to infect 293T-hACE2 cells in presence of increasing concentrations of EH3 (**C**) or EH8 (**D**) at 37°C for 1 h prior infection of 293T-ACE2 cells. (**E**) Cell-to-cell fusion was measured between 293T effector cells expressing HIV-1 Tat and SARS-CoV-2 S D614G which were incubated in presence of increasing concentrations of CV3-1, EH3 or EH8 at 37 °C for 1 h prior coculture with TZM-bl-hACE2 target cells. (**C-E**) Fitted curves and IC50 values were determined using a normalized nonlinear regression. (**F**) Cell-surface staining of CEM.NKr-Spike (Wuhan-Hu-1 strain) using increasing concentrations of CV3-1, EH3 or EH8 mAbs. (**G**) Parental CEM.NKr cells were mixed at a 1:1 ratio with CEM.NKr-Spike cells and were used as target cells. Cryopreserved PBMCs from uninfected donors were used as effector cells in a fluorescence-activated cell sorting (FACS)-based ADCC assay. The graphs shown represent the percentages of ADCC obtained in the presence of increasing concentrations of CV3-1, EH3 or EH8 mAbs. These results were obtained in at least 3 independent experiments. Error bars indicate mean ± standard error.

To evaluate the affinity between EH3 or EH8 with the different VOCs and VOIs, we performed surface plasma resonance (SPR) measurements using recombinant SARS-CoV-2 RBDs. Antibodies were separately immobilized on Protein-A sensor chips and RBDs of WT and 6 VOCs were used as the flow analytes (**Figures S2A-B, Table 1**). EH3 and EH8 had similar low nanomolar affinity to the WT RBD (equilibrium dissociation constants (KD) in the range of 0.83-2.79 nM and 1.30-7.22 nM, respectively). However, their kinetic features were distinct: EH3 had a 3.1-4.8-fold slower association constant or ‘on-rate’ (*kon* from 8.32 × 10^4^ to 2.74 × 10^5^ M^-1^s^-1^) than EH8 (*kon* from 4.03 × 10^5^ to 8.37 × 10^5^ M^-1^s^-1^) but also displayed a slower dissociation constant or ‘off-rate’, with EH8 having a *k*off 4.8-12.5 -fold faster than EH3. The much faster off-rates for EH8 suggest that it either forms a less stable antigen complex than EH3 or that its epitope footprint is smaller than EH3. Both antibodies maintained strong low nanomolar binding affinities towards all tested RBD variants except for RBDB.1.1.529.1 (Omicron, BA.1) (**Figures S2A-B, Table 1**). Indeed, both EH3 and EH8 displayed a dramatic reduction in KD to RBDB.1.1.529.1 (842- and 475- fold, respectively) with 10- to 18-fold decrease in on-rates (*k*on) and 26- to 78-fold increase in off- rates (*k*off) (**Table 1**). These results are in line with the impaired neutralization activity of both antibodies against the B.1.1.529 variants with IC50 values mostly beyond the range of detection (**Figure 2D**).

**Table 1.** Summary of SPR kinetic constants.

| Immobilized ligand | Flow analyte | Fitting mode | $k_{on}$ ($M^{-1} s^{-1}$) | $k_{off}$ ($s^{-1}$) | $K_D$ (nM) | Chi<sup>2</sup> value<sup>#</sup> | $K_D$ reduction fold compared to RBD<sub>WT</sub> |
| --- | --- | --- | --- | --- | --- | --- | --- |
| EH3 IgG | RBD<sub>WT</sub><sup>*</sup> | 1:1 | $8.32 \times 10^4$ | $2.32 \times 10^{-4}$ | 2.79 | 1.18 | 1 |
| | RBD<sub>WT</sub> | 1:1 | $2.74 \times 10^5$ | $2.28 \times 10^{-4}$ | 0.83 | 1.05 | 1 |
| | RBD (B.1.1.7, Alpha) | 1:1 | $7.58 \times 10^4$ | $2.28 \times 10^{-3}$ | 17.8 | 2.76 | 21.4 |
| | RBD* (B.1.351, Beta)* | 1:1 | $5.81 \times 10^4$ | $3.23 \times 10^{-3}$ | 55.5 | 1.08 | 19.9 |
| | RBD (B.1.429, Epsilon) | 1:1 | $1.58 \times 10^5$ | $1.44 \times 10^{-4}$ | 0.92 | 0.42 | 1.11 |
| | RBD (B.1.526, Iota) | 1:1 | $1.78 \times 10^5$ | $2.33 \times 10^{-4}$ | 1.31 | 0.21 | 1.57 |
| | RBD (B.1.617.2 Delta) | 1:1 | $1.02 \times 10^5$ | $1.29 \times 10^{-3}$ | 12.7 | 1.51 | 15.3 |
| | RBD (B.1.1.529.1, Omicron BA.1) | 1:1 | $2.56 \times 10^4$ | $1.79 \times 10^{-2}$ | 699 | 1.89 | 842 |
| EH8 IgG | RBD<sub>WT</sub><sup>*</sup> | 1:1 | $4.03 \times 10^5$ | $2.91 \times 10^{-3}$ | 7.22 | 1.54 | 1 |
| | RBD<sub>WT</sub> | 1:1 | $8.37 \times 10^5$ | $1.09 \times 10^{-3}$ | 1.30 | 2.12 | 1 |
| | RBD (B.1.1.7, Alpha) | 1:1 | $8.81 \times 10^4$ | $2.07 \times 10^{-3}$ | 23.5 | 2.65 | 18.1 |
| | RBD* (B.1.351, Beta)* | 1:1 | $1.27 \times 10^5$ | $2.86 \times 10^{-3}$ | 22.5 | 1.76 | 3.1 |
| | RBD (B.1.429, Epsilon) | 1:1 | $7.32 \times 10^5$ | $1.13 \times 10^{-3}$ | 1.54 | 1.28 | 1.2 |
| | RBD (B.1.526, Iota) | 1:1 | $4.08 \times 10^5$ | $1.50 \times 10^{-3}$ | 3.67 | 0.64 | 2.82 |
| | RBD (B.1.617.2 Delta) | 1:1 | $6.54 \times 10^4$ | $2.46 \times 10^{-3}$ | 37.6 | 0.48 | 28.9 |
| | RBD (B.1.1.529.1, Omicron BA.1) | 1:1 | $4.57 \times 10^4$ | $2.88 \times 10^{-2}$ | 618 | 2.07 | 475 |

\* Minimum RBD segment (residue 329-527). Otherwise, the rest are residue 319-537.

### Molecular Basis for EH3 and EH8 binding to SARS-CoV-2 RBD

To better understand the differences in binding affinities and neutralization potency of EH3 and EH8, we solved the crystal structures of their antigen binding fragments (Fabs) with the SARS-CoV-2 WT RBD at 2.65 Å and 2.49 Å resolution, respectively (**Figures 3-4** and **Table S2**). Both antibodies rely on their heavy chains to interact with the RBD, with a total buried surface area (BSA) of 1069.7 Å^2^ for EH3 and 632.0 Å^2^ for EH8 (**Figure 4D**). We observed that EH3 and EH8 bind to the RBD from distinct orientations and recognize different subsets of RBD residues (**Figures 3A-C**), partly spanning the receptor binding motif (RBM). As shown in **Fig. 3A-C**, the EH3 epitope on the RBD extensively overlaps with the ACE2 binding site. EH8, on the other hand, only interacts with the protruding RBD ridge. As a typical IGHV3-53 encoded antibody, EH3 adopts a common RBD binding mode that utilizes germline residues within CDR H1, CDR H2 and a short CDR H3 (10mer) that covers the majority of RBM. EH3 engages 32 RBD residues, 19 of which are involved in ACE2 binding. In comparison, EH8 only interacts with 14 RBD residues, all of which are in the exposed and mobile ridge region (residues 475-481 and 483-489). Our epitope analysis indicates that both EH3 and EH8 can block the ACE2-RBD interaction and function as competitive inhibitors (**Figures 3B-C**). Structural superimposition of both Fabs onto the SARS-CoV-2 Spike trimer in the “one-RBD-up” and fully closed (all-RBD-down) conformations revealed that epitope of EH3 is inaccessible in the closed Spike conformation. Therefore, it is only able to bind to RBD in the ‘up’ conformation (**Figure 4B**). In contrast, EH8 can engage the RBD ridge in both conformations (**Figure 4C**). EH3 and EH8 can therefore be assigned to Class-1 and Class-2 RBD-targeting antibodies, respectively, in accordance with the categories defined by Barnes *et al.* ^26^.

**Figure 3.**
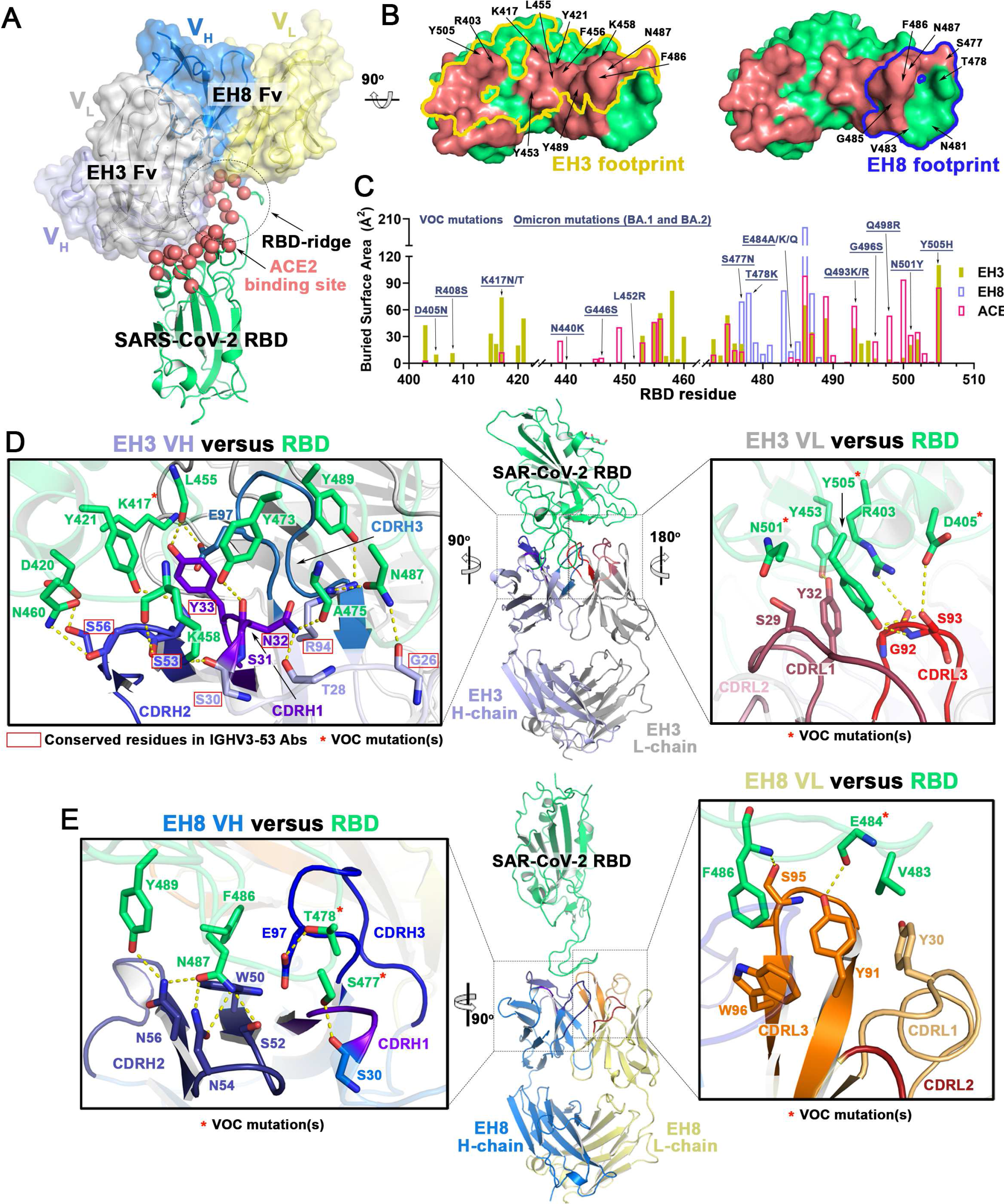
Molecular basis of SARS-CoV-2 S recognition by EH3 and EH8 antibodies. (**A**) Crystal structures of EH3 Fab-RBD and EH8 Fab-RBD complexes. Structures were superimposed based upon the RBD domain and are shown as ribbons. The molecular surfaces are displayed over the variable regions of both Fabs (constant domains were omitted for simplicity). The Cα atoms of the RBD that contribute to receptor ACE2 binding residues are highlighted as red spheres**. (B)** Epitope footprints of EH3 (yellow line) and EH8 (blue line) contoured over the RBD surface (green). ACE2 binding sites are colored in red and the major RBD residues known to contribute to Fab-RBD interactions are labeled. **(C)** Diagram of the buried surface area (BSA) of RBD residues contributing to EH3-RBD (yellow), EH8-RBD (blue) and ACE2-RBD (red) interfaces. The BSA of individual RBD residues was calculated using PISA ^80^ and is given as the sum contributed by the heavy and light chains of the antibody or as the average contributed by ACE2 in two high-resolution structures (PDB IDs 6VW1 ^81^ and 6M0J ^82^). Residue mutations found in VOCs are denoted above with those occurring in the Omicron variant underlined. **(D-E**) The interaction network at the (**D**) EH3-RBD and (**E**) EH8-RBD interface. The Fab-RBD structures are shown in the center with close-up views showing contacts mediated by the antibody’s CDRs to the RBD (green). Hydrogen bonds and salt bridges with bond lengths < 3.5 Å are depicted as yellow dashed lines and the VOCs mutations are marked with red asterisks. IGHV3-53 conserved residues in **D** are highlighted by red boxes.

**Figure 4.**
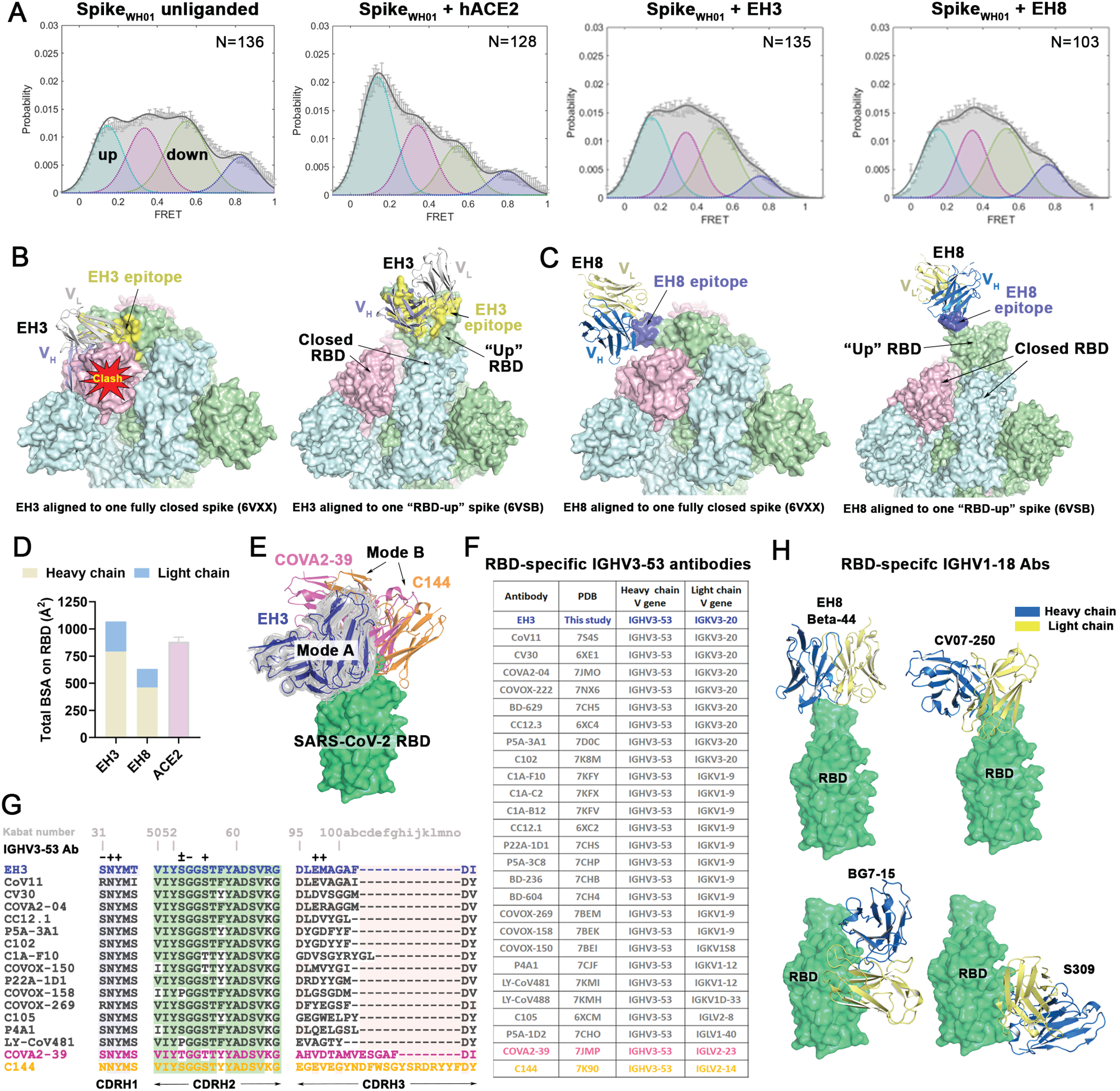
Epitope-driven S conformational preference by RBM-binding EH3 and Ridge- directed EH8. (**A**) Conformational states of SWT on VLPs surface monitored by smFRET in ligand-free state, or in the presence of ACE2, EH3 IgG or EH8 IgG. FRET histograms with the number (N) of individual dynamic molecules/traces compiled into a conformation-population FRET histogram (gray lines) and fitted into a 4-state Gaussian distribution (solid black) centered at 0.1 FRET (dashed cyan), 0.3 FRET (dashed red), 0.5 FRET (dashed green), and 0.8 FRET (dashed magenta). (**B-C**) EH8 but not EH3 recognized both RBD-up and RBD-down S. Structural superimposition of (**B**) EH3 (left panel) and (**C**) EH8 (right panel) onto SARS-CoV-2 Spike in fully closed conformation (6VXX ^83^, left) and “one-RBD-up” conformation (6VSB ^84^, right). Epitopes of EH3 and EH8 are colored in yellow and blue, respectively. Structural alignments show potential clashes for the heavy chain of EH3 with the adjacent RBD when bound to the closed Spike, while the ridge-binding EH8 can bind to both the RBD-up and RBD-down Spike with no steric clashes. (**D**) The total buried surface area (BSA) at the Fab-RBD interface contributed by heavy/light chains of two nAbs and ACE2 (average of two ACE2-RBD crystal structures 6M0J ^82^ and 6VW1 ^81^). (**E-F**) EH3 in context of other IGHV3-53 antibodies. (**E**) Structures of EH3-RBD and the IGHV3-53 antibodies available in PDB in complexes with SARS-Cov-2 antigen were superimposed based upon the RBD. The constant regions of the Fabs were omitted for clarity. (**F**) The table (right panel) shows a list of IGHV3-53 antibodies used for the structural alignment. EH3 is shown (blue) superimposed on the 26 structurally available IGHV3-53 derived antibodies (semi-transparent grey ribbons) that bind with similar angle. Two antibodies, COVA2-39 and C144, which bind using different angles of approach, are colored pink and orange respectively. (**G**) Heavy chain CDRs sequence alignments of EH3 with 16 selected RBD-specific IGHV3-53 antibodies. The identical amino acids as compared to EH3 are shaded in colors. Residues involved in salt-bridges or H-bonds to the RBD are marked above the sequence with (+) for the side chain, (-) for the main chain and (±) for both. (**H**) Comparison of four structurally resolved RBD-directed IGHV1-18 antibodies, including EH8, Beta-44 (7PS6) ^42^, BG7-15 (7M6G) ^40^, CV07-250 (6XKQ) ^85^ and S309 (6WPT) ^41^, showing distinct RBD epitopes and angles of approach. The constant regions of the Fabs were omitted for clarity.

To confirm these interpretations, we used single-molecule fluorescence resonance energy transfer (smFRET) imaging to characterize the dynamic conformational preferences of EH3 and EH8 bound to the prototypic Wuhan-Hu-1 Spike on the surface of virus-like particles (VLPs), as previously described ^24^. In its unliganded form, the RBD-down ground state was the dominant S conformation. The addition of the receptor ACE2 induced the S transition from the RBD-down to the RBD-up state (**Figure 4A**). As expected, EH3 acted like other RBD-specific antibodies in mimicking receptor ACE2 and shifted the S conformational equilibrium from the ∼0.5 FRET RBD-down state to the ∼0.1 FRET RBD-up state, albeit to a lower extent than ACE2. In agreement with its ability to recognize the RBD-down and RBD-up conformations, the Spike conformational landscape remained relatively unchanged in the presence of EH8, confirming that this antibody does not display S conformational preference (**Figures 4A, C**). Interestingly, there seems to be a correlate between the ability of RBD-specific antibodies to stabilize the RBD-up conformation and their ability to induce S1 shedding, and this phenomenon appears to be epitope-dependant (**Figure S3**). Class 1 antibodies such as EH3, CV3-1 and Casirivimab induce potent S1 shedding and are known to stabilize the RBD-up conformation. Class 2 antibodies such as EH8 induce modest S1 shedding and are conformational-independent. Finally, class 3 antibodies such as Imdevimab do not induce S1 shedding (**Figure S3**).

### EH3 acts like a canonical IGHV3-53 NAb

The molecular interactions of EH3-RBD highly resembles those in the reported crystal structures of SARS-CoV-2 RBD and IGHV3-53 antibodies with short CDR H3s. We identified 26 IGHV3- 53 gene-derived RBD binding antibodies with available crystal or Cryo-EM structures from the CoV3D database^27^ (**Figures 4F, S4A**). Most exhibited little affinity maturation with high neutralization potency ^8, 9, 26, 28–38^. Despite of the diverse use of light chain V genes, 24 out of 26 of these antibodies whose CDR H3 are 7-13 residues in length, share a very similar angle of approach to the RBD and make contact to similar residues on the RBD (mode A). Two exceptions, COVA2- 39 ^36^ and C144 ^26^, which have significantly longer CDR H3 loops (15mer and 23mer respectively), bind to RBD from very different orientations (mode B, **Figure 4E**). This CDR H3-governed binding mode in IGHV3-53 encoded antibodies has been previously discussed by Meng Y. *et al* ^8^ and Wu N. C. *et al* ^36^. The limited space within the paratope-epitope interface in binding mode A can only be accommodated by a short CDR H3. In contrast, in binding mode B there are fewer structural constraints for CDR H3 length. Of note, C144 of the latter group is the only IGHV3-53 encoded antibody that can recognize RBD in its down conformation ^26^.

The germline-encoded CDR H1 and CDR H2 of EH3 have highly conserved sequences and identical lengths (5-mer and 16-mer, respectively) which is like the other RBD specific IGHV3-53 encoded antibodies identified to date (**Figures 4G, S4A**). There are 19 hydrogen bonds and 1 salt-bridge found in the EH3-RBD interface, 12 of which involve the side chains of EH3 residues, including N32 and Y33 from CDR H1, S53 and S56 from CDR H2, E97 and M98 from CDR H3, and T28 and R94 from framework regions 1 and 3 (FR1 and FR3). Site-directed mutagenesis studies showed that, although EH3 interacts with an extended RBM epitope, its interaction with RBD is highly dependent on a network of aromatic residues involving two invariant interactions, S53CDRH2-Y421RBD and Y33CDRH1-F456RBD, which appear to be critical for EH3 recognition (**Figure 5A**), consistent with the high BSA contributed by these two aromatic RBD residues (**Figure 3C**). F456RBD has also been found to be critical for ACE2 interaction (**Figure 5C**), suggesting that mutations abrogating EH3 binding would result in a high fitness cost for the virus, which would require compensating mutations elsewhere in the RBM to retain ACE2 binding. The highly conserved RBD contacts with the EH3 residues from the framework as well as CDR H1 & H2 underlie the general structural basis for IGHV3-53 NAbs’ broad neutralization against SARS-CoV-2 VOCs.

**Figure 5.**
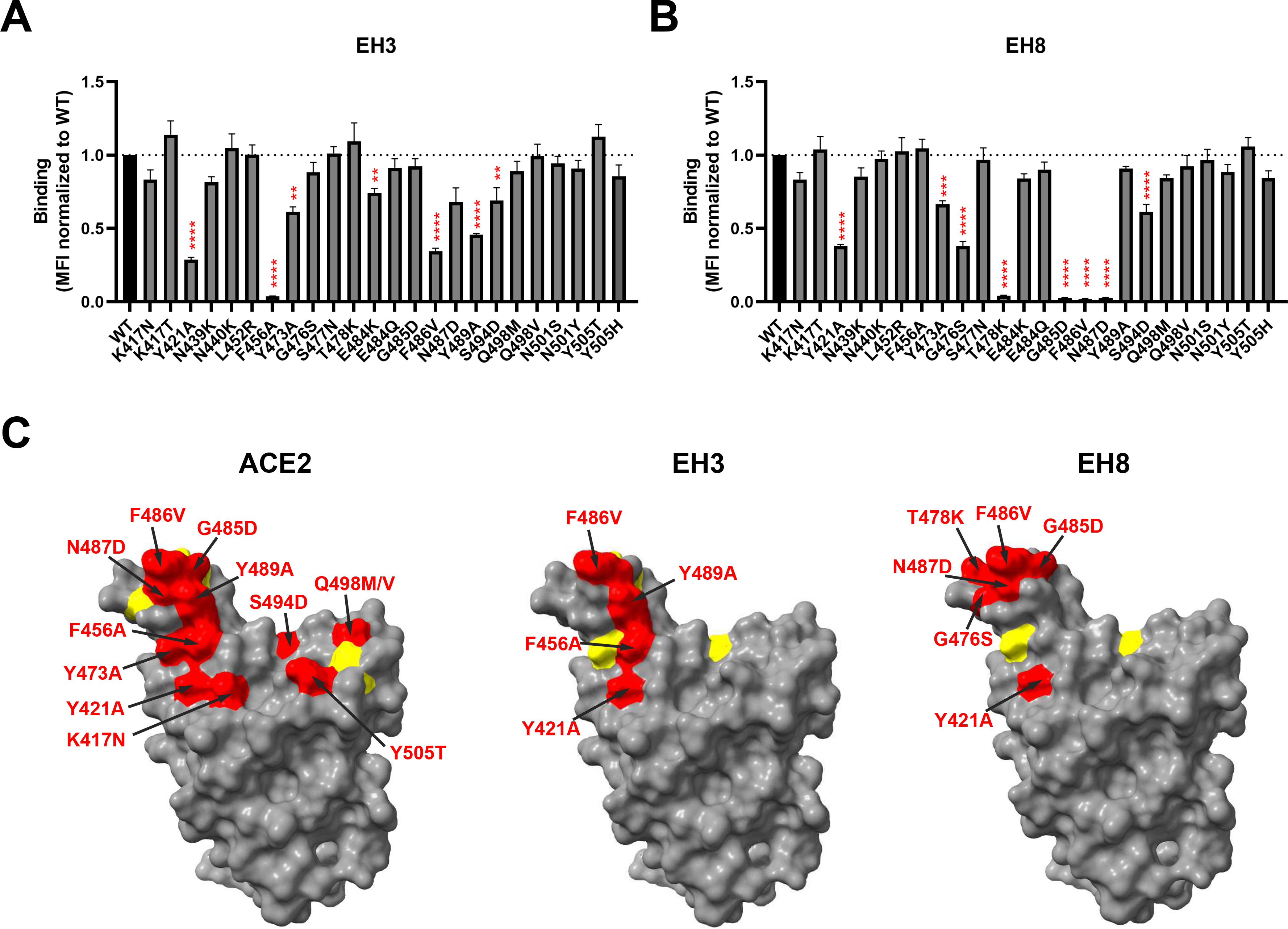
Epitope mapping of RBD-specific mAbs by site-directed mutagenesis. (**A-B**) Cell-surface staining of 293T cells expressing selected full-length SARS-CoV-2 S harboring RBM mutations using EH3 (**A**) and EH8 (**B**). The graphs shown represent the median fluorescence intensities (MFI) corrected for cell-surface S expression of the corresponding mutant using the CV3-25 mAb and further normalized to the MFI obtained with S D614G (WT). Dashed lines indicate the reference value obtained with S D614G (WT). Error bars indicate means ± SEM. These results were obtained in at least three independent experiments. Statistical significance was tested using one-way ANOVA with a Dunnett post-test (∗p <0.05; ∗∗p <0.01; ∗∗∗p <0.001; ∗∗∗∗p <0.0001). (**C**) Structural representation of SARS-CoV-2 RBD depicted as a surface model (PDB: 6VW1) using UCSF ChimeraX. Amino acid substitutions able to significantly decrease the binding of indicated ligands are colored in red (if decrease is more than 50% compared to WT) or in yellow (if decrease is less than 50% compared to WT).

The light chain of EH3 contributes significantly less to the RBD interface, reflected in its low BSA (278.2 Å^2^) as compared to the heavy chain (791.5 Å^2^). Indeed, use of light chain V genes is more variable among the IGHV3-53 encoded RBD specific antibodies, with two Kappa light chains, IGKV3-20 and IGKV1-9, occurring most often (**Figure 4G**). EH3 utilizes the IGKV3-20 germline sequence and interacts with the flat surface of the RBD core through CDR L1 and CDR L3 (**Figure 3D**). Two VOCs mutations are located within the light chain-RBD interface, N501Y and Y505H, but they do not seem to impact significantly EH3 binding, confirming that the heavy chain-RBD interface is the major contributor for its interaction with Spike (**Figure 5A**).

Notably, except for the recent Omicron BA.1 variant, the RBD binding and neutralization potency for EH3 against the tested VOCs were only reduced by 1.5- to 20-fold and 1.1- to 2.6-fold respectively (**Figures 1D, 2D, S2A and Table 1**). Our structural and mutational analysis illustrated that a combination of mutations within the paratope-epitope interface might be necessary to impact EH3 binding, including K417N/T, S477N, F486V, Q493K/R, N501Y and Y505H (**Figures 3D and 5A**), while biolayer interferometry analyses indicated that E484K, located at the periphery of the EH3 epitope, might exert only mild effect on EH3 binding (**Figure S2C**).

### EH8 recognizes a novel RBD-ridge epitope that is exposed in both the RBD-down and RBD- up Spike conformations

The ridge-binding EH8 is encoded by rarely used V genes for SARS-CoV-2 antibodies, IGHV1- 18 and IGLV-2-23 ^31^. EH8 has an epitope footprint of 632.0 Å^2^ which makes it one of the smallest epitope footprints according to an excellent review summarizing 38 RBD-specific mAbs with known structures ^39^. EH8 engages a minimum set of RBD ridge residues (475-489) and has a substantially lower degree of overlap with ACE2 as compared to other RBM-binding mAbs (**Figures 3A-C**) such as EH3. Despite the narrow epitope, its RBD binding affinity (**Figures 2A-B & S2A-B**, **Table 1**) and neutralization potency (**Figures 2C-D**) against the early VOCs are comparable to EH3. The EH8-RBD interface consists of 9 hydrogen bonds contributed mainly by CDR H2, CDR H3 and CDR L3 (**Figure 3E**). The ridge residue F486RBD, which is conserved among all the earlier SARS-CoV-2 VOCs (except the most recent BA.4 and BA.5) and is one of the most targeted residues by RBD-binding mAbs, has the highest single residue BSA (203.9 Å^2^ out of the total 632.0 Å^2^ BSA of the interface). The phenyl side chain of F486RBD inserts into a hydrophobic pocket in EH8 formed by W50CDRH2 and S95-W96CDRL3 (**Figure 3E**). In addition, N487RBD forms a hydrogen bonding network with three CDR H2 residues, S52, N54 and N56. T478RBD forms a hydrogen bond with E97 on CDR H3 (**Figure 3E**). Using a panel of RBM mutants, we were able to confirm the importance of T478, F486 and N487 for EH8 interaction with the RBD ridge (**Figure 5B**). Our results suggest that the T478K mutation that arose in the B.1.617.2 (Delta) and B.1.1.529 (Omicron) lineages is sufficient to confer resistance to EH8 binding (**Figure 5B**), in line with the markedly reduced neutralization by EH8 against these VOCs (**Figure 2D**).

Although recent studies have identified several IGHV1-18 antibodies (e.g. BG7-15 ^40^ and S309 ^41^) that are able to bind the closed Spike, they recognize different epitopes (**Figure 4H**). To our knowledge, EH8 and a recently reported Beta-44^42^ are the only antibodies that solely interact with the RBD ridge. It can bind in an upright orientation from the top in an open Spike trimer or in a nearly perpendicular orientation to the trimer axis in a closed Spike (**Figure 4C**). Sequence analysis of CDR residues suggests that the high diversity in the angle of approach to the RBD for IGHV1-18 encoded antibodies is likely dictated by the length and sequence of CDR H3 (**Figure S4B**) which has uniformly undergone a high degree of somatic hypermutation to acquire RBD binding affinity.

### EH3 effectively protects K18-hACE2 mice from lethal infection of the Delta VOC

We evaluated *in vivo* efficacy of EH3 against Delta VOC in the stringent K18-hACE2 mouse model. K18-hACE2 mice were administered EH3 or hIgG1, 12 h before lethal challenge with reporter SARS-CoV-2 Delta-nLuc VOC to enable whole-body bioluminescence imaging (BLI) (**Figure 6A**). Quantification of nLuc signals (flux) following longitudinal non-invasive BLI and terminal imaging of target organs revealed that compared to hIgG1-treated mice, prophylactic administration of EH3 efficiently inhibited Delta-nLuc VOC replication in the lungs, nose and prevented neuroinvasion. (**Figures 6B-D, G, H**). In contrast to mice in control cohorts that steadily lost body weight, EH3-pretreated mice did not experience weight loss (**Figure 6E**). Accordingly, 100% of the EH3-treated mice were protected from Delta-nLuc-induced mortality. (**Figure 6F**). An effective virologic control and reduced pathogenesis was also demonstrated by significantly reduced viral loads as well as inflammatory cytokines in target organs (nose, lung and brain) of mice under EH3 prophylaxis in comparison with control mice. (**Figures 6I-L**). Overall, these data demonstrate that EH3, as the representative of the prevalent IGHV3-53 encoded RBD-specific antibodies, remains highly effective in controlling the SARS-CoV-2 Delta VOC.

**Figure 6.**
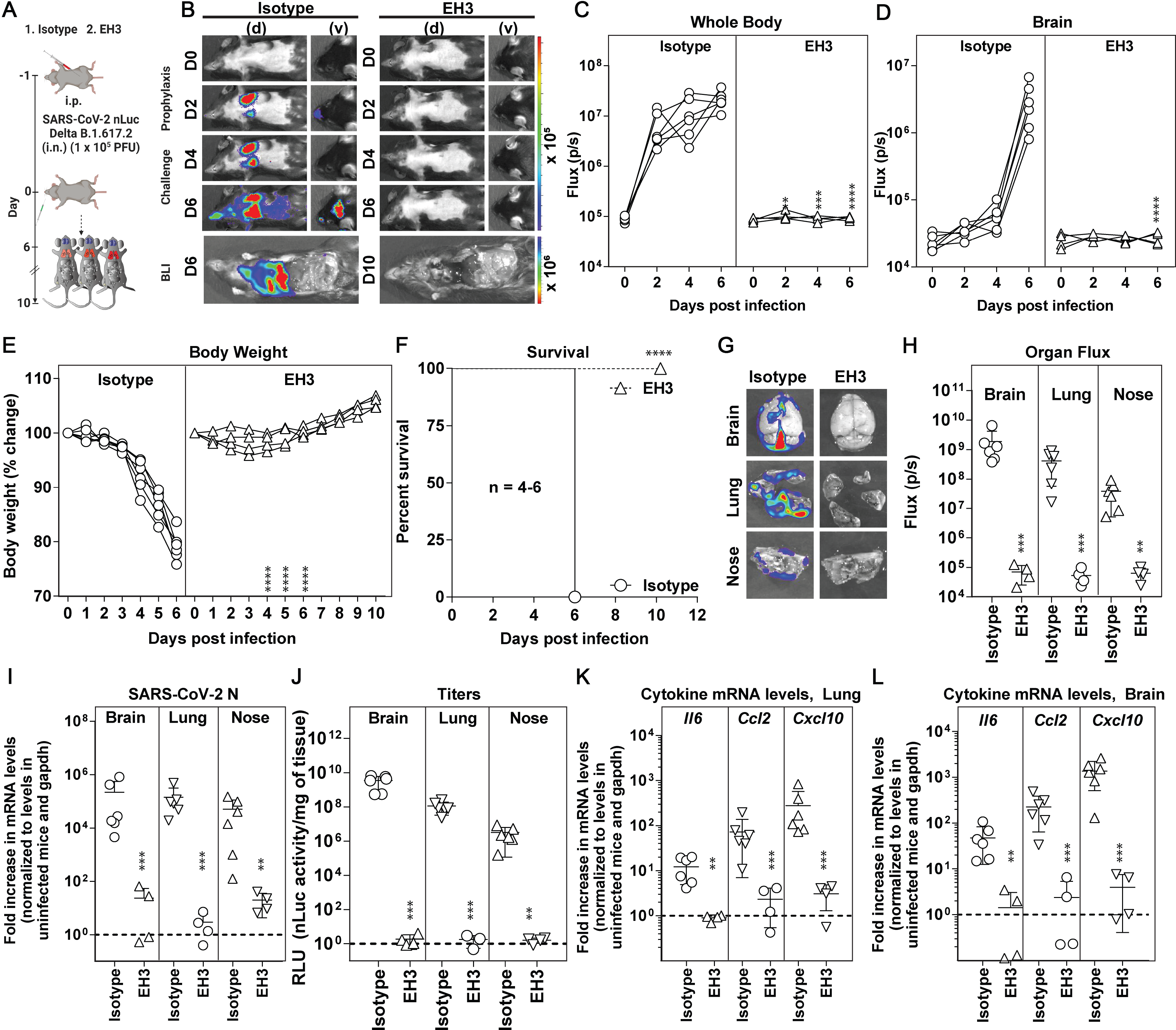
Prophylactic administration of EH3 effectively protected K18-hACE2 mice from lethal challenge of SARS-CoV-2 Delta variant. (**A**) Schematic presenting the experimental design for testing *in vivo* efficacy of the EH3 antibody delivered at 12.5 mg/kg body weight intraperitoneally (i.p.) in a prophylactic regimen 12 h before challenging K18-hACE2 mice with 1 × 10^5^ PFU SARS-CoV-2 Delta-nLuc VOC. Human IgG1- treated (n=6) mice were used as control. (**B**) Representative BLI images of SARS-CoV-2-nLuc- infected mice in ventral (v) and dorsal (d) positions. (**C-D**) Temporal quantification of nLuc signal as flux (photons/sec) computed non-invasively. (**E**) Temporal changes in mouse body weight with initial body weight set to 100% for an experiment shown in A. (**F**) Kaplan-Meier survival curves of mice (n = 4-6 per group) statistically compared by log-rank (Mantel-Cox) test as in A. (**G-H**) *Ex vivo* imaging of indicated organs and quantification of nLuc signal as flux (photons/sec) after necropsy as per A. (**I**) Fold changes in nucleocapsid mRNA expression in brain, lung, and nasal cavity tissues. Data were normalized to Gapdh mRNA in the same sample and in non-infected mice after necropsy. (**J**) Viral loads (nLuc activity/mg) from indicated tissues using Vero E6 cells as targets. Virus loads in indicated tissues were determined when they succumbed to infection and at 10 days post-infection (dpi) for surviving mice. (**K-L**) Fold-change of cytokine mRNA expression in brain and lung tissues. Data were normalized to Gapdh mRNA in the same sample and in non-infected mice after necropsy. Cytokines in indicated tissues were determined when they succumbed to infection and at 10 dpi for surviving mice. Grouped data in (C-E) were analyzed by two-way ANOVA followed by Tukey’s multiple comparison tests. The data in (H-L) were analyzed by t test followed by non-parametric Mann-Whitney U tests. ∗, p < 0.05; ∗∗, p < 0.01; ∗∗∗, p < 0.001; ∗∗∗∗, p < 0.0001; Mean values ± SD are depicted.

## DISCUSSION

Understanding how antibodies are elicited in different population to combat SARS-CoV-2 infection, how they contribute to disease severity, and how escape mutations emerge in response to antibody selection, could aid in the development of safe, effective SARS-CoV-2 therapeutics and vaccines. In this study, we isolated two potently neutralizing mAbs, EH3 and EH8, from a 6- year-old patient suffering from MIS-C during the early onset of the COVID-19 pandemic. Both Abs potently neutralized SARS-CoV-2 Wuhan-Hu-1 strain, D614G, B.1.1.7, B.1.351 and P.1 with IC50 < 0.04 μg/mL and mediated robust Fc-dependent elimination of S-expressing cells. EH3 remained highly effective against B.1.617.2 (Delta) *in vitro* and *in vivo,* however this variant was resistant to EH8. The binding potency of both NAbs to Omicron BA.1 RBD decreased by 400- to 800-fold. High-resolution crystal structures, together with mutagenesis studies and smFRET, provide the molecular basis of epitope-driven neutralization, Fc-effector functions, and VOCs evasion for both Abs.

To date, IGHV3-53 has been found to be the most frequently used IGHV gene among the effective SARS-CoV-2 RBD-specific antibodies ^8, 9^. EH3 is a representative IGHV3-53-encoded NAb which possesses a short CDR H3 with low somatic hypermutation (SHM), shared structural features and broad reactivity against multiple SARS-CoV-2 VOCs prior to the emergence of the Omicron variant. Our crystal structure of EH3-RBD complex revealed that the well-conserved CDR H1 and CDR H2 (**Figure 4G**) establish extensive hydrogen bonding networks with the invariant RBM residues (e.g. Y421, Y473, N487, etc.), which likely underlie the broad neutralization breadth of IGHV3-53 NAbs ^10^. Of note, the high variation in the CDR H3 sequence and length among IGHV3-53 Abs not only alters the loop conformations but also dictates the RBD binding angles ^36^. Our *in vivo* prophylactic studies on the susceptible K18-hACE2 mouse model demonstrated that a single administration of EH3 conferred 100% protection from the lethal challenge by the SARS-CoV-2 Delta variant, providing a rationale for the prevalence of IGHV3- 53 antibodies–elicited by natural infection and/or vaccination as they likely remain effective against many SARS-CoV-2 VOCs.

Another pediatric NAb EH8, represents a minority class of IGHV1-18 encoded antibodies that can recognize both the RBD-down and RBD-down Spike conformations. EH8 features a narrow RBD-ridge epitope (residue 475-489) where a diversity of immune-evading mutations (e.g., S477N, T478K, E484A/K/Q and F486V) have emerged in SARS-CoV-2 VOCs. As a result, the neutralization breadth of EH8 is substantially limited, particularly to the most recent VOCs (Delta, Omicron BA.1 and BA.2). Nevertheless, we hypothesize that the potent ADCC activity mediated by EH8 is due to its ability to indiscriminately recognize the S trimers on infected cells, regardless of the RBD conformations. While it has been previously suggested that Ab-based interventions capable of robust Fc-effector functions could exacerbate COVID-19 severity by inducing adverse cytokine cascades and enhancing macrophage infections, several *in vivo* studies have demonstrated the absence of antibody-dependent enhancement (ADE) of infection^43^ and the beneficiary contribution from Fc-dependent antiviral activities in prophylactic and therapeutic settings^19–21, 44^.

The S antigenicity, affinity to hACE2 and susceptibility to RBD-specific NAbs elicited by the existing VOCs and ancestral-S based vaccines are dramatically altered by the accumulation of >30 total mutations, 14–16 of which reside in the RBD, in Omicron variants as opposed to the earlier VOCs, which only contained up to 3 RBD mutations. We found that the T478K found in Delta and Omicron variants is sufficient to evade the binding of EH8. The reduced binding and neutralization potency of EH3 to Omicron may be an accumulative effect of several moderately affecting mutations, including K417N, S477N, E484A, Q493R, N501Y and Y505H. An Omicron-specific booster shot is urgently needed to control these widespread variants. The recent emergence of Omicron BA.4 and BA.5 lineages, derived from the BA.2 lineage, could lead to further antibody escape due to the presence of additional mutations in the RBM, such as L452R and F486V^45–48^. Given the high variability of the RBM, development of vaccines or immunotherapies targeting more conserved epitopes such as the S2 stem helix^25, 49–55^ or fusion peptide^56, 57^ should be considered.

Taken together, our data provides a comprehensive functional, biophysical, structural, and *in vivo* study of two pediatric antibodies elicited by natural infection with a nascent SARS-CoV-2 strain. Detailed epitope mapping and mutational analysis interpreted the cross-reactivity of both Abs against the early SARS-CoV-2 VOCs as well as the high resistance of recent Omicron variants. This study also provides a blueprint for understanding the correlates between antibody function and structure, viral mutation frequencies under Abs selection and design of potential Spike/RBD- based immunogens for vaccines to mitigate future pandemics.

## ACKNOWLEDGEMENTS

The authors thank the CRCHUM BSL3 and Flow Cytometry Platforms for technical assistance. We thank Dr Stefan Pöhlmann and Dr Markus Hoffmann (Georg-August University) for the plasmids coding for SARS-CoV-2 Spike Wuhan-Hu-1 and Dr M. Gordon Joyce (U.S. MHRP) for the CR3022 monoclonal antibody. CV3-1 and CV3-25 antibodies were produced using the pTT vector kindly provided by the Canada Research Council. Use of the Stanford Synchrotron Radiation Lightsource, SLAC National Accelerator Laboratory, is supported by the U.S. Department of Energy, Office of Science, Office of Basic Energy Sciences under Contract No. DE-AC02-76SF00515. The SSRL Structural Molecular Biology Program is supported by the DOE Office of Biological and Environmental Research, and by the National Institutes of Health, National Institute of General Medical Sciences. We thank the staff members of the SER-CAT and GM/CA beamlines at the Advanced Photon Source, Argonne National Laboratory for beamline support.

This work was supported by a Canadian Institutes of Health Research (CIHR) operating grant Pandemic and Health Emergencies Research/Project #465175 to M.P., W.M., and A.F., by NIH grant R01AI163395 to WM, by Uniformed Services University of the Health Sciences (USUHS) intramural funds to M.P., by le Ministère de l’Économie et de l’Innovation (MEI) du Québec, Programme de soutien aux organismes de recherche et d’innovation to A.F., the Fondation du CHUM, a CIHR foundation grant #352417 to A.F., CIHR stream 1 and 2 for SARS-CoV-2 Variant Research to A.F. and M.C., an Exceptional Fund COVID-19 from the Canada Foundation for Innovation (CFI) #41027 to A.F., the Sentinelle COVID Quebec network led by the Laboratoire de Santé Publique du Quebec (LSPQ) in collaboration with Fonds de Recherche du Québec-Santé (FRQS) and Genome Canada—Génome Québec, and by the Ministère de la Santé et des Services Sociaux (MSSS) and MEI to A.F, and by the Fonds Ligne from the Fondation de l’Hôpital Sainte-Justine to E.H, the Banque de Montreal (BMO) Pediatric Pediatric Immunology Research Chair to E.H. A.F. and M.C. are recipients of Canada Research Chairs on Retroviral Entry no. RCHS0235 950232424 and CRC in Molecular Virology and Antiviral Therapeutics, respectively. M.A.S is partially supported by a FRQS Junior 1 fellowship. J.P. and S.P.A. are supported by CIHR doctoral fellowships, G.B.-B. by a FRQS doctoral fellowship, M.W.G. by the Gruber Science fellowship and R.G. by a MITACS Accélération postdoctoral fellowship. The funders had no role in study design, data collection and analysis, decision to publish, or preparation of the manuscript.

## Author Contribution

**Conceptualization:** Y.C., J.P., I.U., M.P., H.R., V.L., V.-P.L., M.A.S., E.H., A.F.; **Methodology:** Y.C., J.P., I.U., H.R., V.L., P.K., B.P., P.D.U., M.A.S., W.M., M.P., A.F.; **Investigation:** Y.C., J.P., I.U., K.B., W.D.T., J.R.G., S.D., S.Y.G., G.B.-B., R.G., M.B., D.V., S.P.A., D.C., G.G., Z.Y., X.B., D. N., R.S., D.W., G.P., D.X., J.R., P.D.U., M.A.S.; **Resources:** M.W.G, Y.B., F.Z., A.R.Z., R.K.H., D.M., M.C.; **Visualization:** Y.C., J.P., K.B., M.A.S; **Formal analysis:** Y.C., J.P.; **Supervision:** P.K., P.D.U., W.M., M.P., E.H., A.F.; **Funding acquisition:** M.C., W.M., M.P., E.H., A.F.; **Writing—original draft:** Y.C., J.P., M.P., K.B., A.F.; **Writing—review and editing:** All authors.

## Disclaimer

The views expressed in this manuscript are those of the authors and do not reflect the official policy or position of the Uniformed Services University, the U.S. Army, the Department of Defense, the National Institutes of Health, Department of Health and Human Services or the U.S. Government, nor does mention of trade names, commercial products, or organizations imply endorsement by the U.S. Government.

## STAR METHODS

## RESOURCE AVAILABILITY

### Lead Contact

Further information and requests for resources and reagents should be directed to and will be fulfilled by the Lead Contact, Andrés Finzi.

### Materials Availability

All unique reagents generated in this study are available from the Lead Contact with a completed Materials Transfer Agreement.

### Data and Code Availability

The published article includes all datasets generated and analyzed for this study. Further information and requests for resources and reagents should be directed to and will be fulfilled by the Lead Contact Author.

## EXPERIMENTAL MODELS AND SUBJECT DETAILS

### Patients and Ethics statement

The patient of interest was enrolled in a cohort of patients with inflammatory and immune disorders, including MIS-C (protocol #3195, CHU Sainte-Justine). PBMCs from healthy individuals (males and females) as a source of effector cells in our ADCC assay were obtained under CRCHUM institutional review board (protocol #19.381). Research adhered to the standards indicated by the Declaration of Helsinki. Written informed consent was obtained for each patient and participant, in accordance with local regulations and with institutional review board approval.

### Cell lines and primary cells

HEK 293T cells (obtained from American Type Culture Collection [ATCC]) were derived from 293 cells, into which the simian virus 40 T-antigen was inserted. 293T-ACE2 cells stably expressing human ACE2 are derived from 293T cells^23^. TZM-bl-ACE2 cells stably expressing human ACE2 are derived from TZM-bl cells and are engineered to contain the Tat-responsive firefly luciferase reporter gene^21^. 293T, 293T-ACE2, and TZM-bl-ACE2 cells were maintained at 37°C under 5% CO2 in DMEM media, supplemented with 5% FBS and 100 U/mL penicillin/ streptomycin. Stably transduced 293T-ACE2 and TZM-bl-ACE2 cells were cultured in medium supplemented with 2 μg/mL of puromycin (Millipore Sigma). CEM.NKr (NIH AIDS Reagent Program) is a T lymphocytic cell line resistant to NK cell-mediated lysis. CEM.NKr-Spike stably expressing SARS-CoV-2 Spike were used as target cells in the ADCC assay^72^. PBMCs were obtained from healthy donor through leukapheresis and were used as effector cells in ADCC assay. CEM.NKr, CEM.NKr-Spike, and peripheral blood mononuclear cells (PBMCs) were maintained at 37°C under 5% CO2 in RPMI media, supplemented with 10% FBS and 100 U/mL penicillin/ streptomycin. Vero E6 (obtained from ATCC) were cultured at 37°C in RPMI supplemented with 10% fetal bovine serum (FBS), 10 mM HEPES pH 7.3, 1 mM sodium pyruvate, 1 × non-essential amino acids, and 100 U/mL of penicillin–streptomycin.

### Viruses

The B.1.617.2 variant (Delta VOC) of SARS-CoV-2 was obtained from BEI resources. We replaced Orf7A with nanoluc luciferase using Circular Polymerase Extension Reaction (CPER)^73, 74^ to obtain luciferase expressing Delta VOC for *in vivo* longitudinal Bioluminescence imaging (BLI) studies. We authenticated Nanoluc reporter Delta VOC by Next-Generation Sequencing at the Yale Center for Genome Analysis using ARTIC primer sets before use in experiments. Viruses were propagated in Vero-E6 cells stably expressing ACE2 and TMPRSS2 by infecting them in T150 cm^2^ flasks at a MOI of 0.1. The culture supernatants were collected after 24 h h when cytopathic effects were clearly visible. The cell debris was removed by centrifugation and filtered through 0.45-micron filter to generate virus stocks. Viruses were then concentrated by adding one volume of cold (4°C) 4x PEG-it Virus Precipitation Solution (40% (w/v) PEG-8000 and 1.2 M NaCl; System Biosciences) to three volumes of virus-containing supernatant. The solution was mixed by inverting the tubes several times and then incubated at 4°C overnight. The precipitated virus was harvested by centrifugation at 1,500 × g for 60 min at 4°C. The concentrated virus was then resuspended in PBS then aliquoted for storage at −80°C. All work with infectious SARS-CoV-2 was performed in Institutional Biosafety Committee approved BSL3 and A-BSL3 facilities at Yale University School of Medicine using appropriate positive pressure air respirators and protective equipment.

### Experimental mouse model

All experiments were approved by the Institutional Animal Care and Use Committees (IACUC) of and Institutional Biosafety Committee of Yale University (IBSCYU). All the animals were housed under specific pathogen-free conditions in the facilities provided and supported by Yale Animal Resources Center (YARC). All IVIS imaging and virus inoculation experiments were done under anesthesia using regulated flow of isoflurane:oxygen mix to minimize pain and discomfort to the animals. Heterozygous transgenic C57BL/6 mice expressing hACE2 under cytokeratin 18 promoter (K18-hACE2) were obtained from the Jackson Laboratory. 6–8-week-old male and female mice were used for all the experiments. The heterozygous mice were crossed and genotyped to select heterozygous mice for experiments by using the primer sets recommended by Jackson Laboratory.

## METHOD DETAILS

### Plasmids

The plasmids expressing SARS-CoV-2 Spike wildtype (Wuhan-Hu-1) and D614G were previously reported^63, 65^. The plasmid encoding for SARS-CoV-2 S RBD (residues 319–541) fused with a hexahistidine tag was previously described^63^. The panel of individual RBM mutations in the full-length SARS-CoV-2 Spike expressor and the Spike from the B.1.429 lineage (S13I, W152C, L452R, D614G) were generated using the QuikChange II XL site-directed mutagenesis kit (Agilent Technologies) and were previously reported^25^. The presence of the desired mutations was determined by automated DNA sequencing. The plasmids encoding the Spike from the B.1.1.7 lineage (Δ69-70, Δ144, N501Y, A570D, D614G, P681H, T716I, S982A and D1118H), the B.1.351 lineage (L18F, D80A, D215G, Δ242–244, R246I, K417N, E484K, N501Y, D614G, A701V), the P.1 lineage (L18F, T20N, P26S, D138Y, R190S, K417T, E484K, N501Y, D614G, H655Y, T1027I) and the B.1.526 lineage (L5F, T95I, D253G, E484K, D614G, A701V) were codon-optimized and synthesized by Genscript^25, 66^. The plasmids encoding the Spike from the B.1.617.1 (E154K, L452R, E484Q, D614G, P681R), the B.1.617.2 (T19R, Δ156–158, L452R, T478K, D614G, P681R, D950N), B.1.1.529.1 (A67V, Δ69-70, T95I, G142D, Δ143-145, Δ211, L212I, ins214EPE, G339D, S371L, S373P, S375F, K417N, N440K, G446S, S477N, T478K, E484A, Q493R, G496S, Q498R, N501Y, Y505H, T547K, D614G, H655Y, N679K, P681H, N764K, D796Y, N856K, Q954H, N969K, L981F) and B.1.1.529.2 (T19I, L24S, Δ25-27, G142D, V213G, G339D, S371F, S373P, S375F, T376A, D405N, R408S, K417N, N440K, S477N, T478K, E484A, Q493R, Q498R, N501Y, Y505H, D614G, H655Y, N679K, P681H, N764K, D796Y, Q954H, N969K) lineages were generated by overlapping PCR using a codon-optimized wild-type SARS-CoV-2 Spike gene that was synthesized (Biobasic, Markham, ON, Canada) and cloned in pCAGGS as a template^25, 67, 68^. All constructs were validated by Sanger sequencing. The plasmid encoding for the ACE2-Fc chimeric protein, a protein composed of an ACE2 ectodomain (1–615) linked to an Fc segment of human IgG1 was previously reported^70^.

### Antibodies and plasma

Plasma from SARS-CoV-2-infected and uninfected donors were collected, heat-inactivated for 1 hour at 56 °C and stored at -80°C until ready to use in subsequent experiments. Plasma from convalescent adult donors S002 (male, 65 years old, 25 days post-symptom onset) and S006 (male, 30 years old, 41 days post-symptom onset) were previously characterized^24^. Plasma from uninfected donors were used as negative controls and used to calculate the seropositivity threshold in the ELISA assay. The monoclonal antibody CR3022 was used as a positive control in ELISA assays and was previously described^58^. The anti-SARS-CoV-2 Spike CV3-1 and CV3-25 mAbs were isolated from the blood of convalescent donor S006 [CV3] using a fluorescent recombinant stabilized Spike ectodomain to probe Spike-specific B cells as previously reported^75^. Horseradish peroxidase (HRP)-conjugated Abs specific for the Fc region of human IgG (Invitrogen) were used as secondary antibodies to detect antibody binding in ELISA experiments. Alexa Fluor-647- conjugated goat anti-human IgG (H+L) Abs (Invitrogen) were used as secondary antibodies to detect antibody binding in flow cytometry experiments. The anti-SARS-CoV-2 RBD rabbit antiserum was used for immunoprecipitation experiments and was previously reported^25^.

### Recombinant protein expression and purification

Recombinant protein expression and purification FreeStyle 293F (ThermoFisher Scientific) cells were grown to a density of 1×10^6^ cells/mL at 37°C with 8% CO2 with regular 135 rpm agitation. Cells were transfected with a plasmid expressing SARS-CoV-2 RBD, a plasmid expressing ACE2-Fc fusion protein or with plasmids expressing the heavy and light chains of a given antibody using ExpiFectamine 293 transfection reagent, as directed by the manufacturer (Thermo Fisher Scientific). One week later, the cells were pelleted and discarded, and supernatants were filtered (0.22-μm-pore-size filter). Recombinant SARS-CoV-2 RBD were purified by nickel affinity columns (Invitrogen) while ACE2-Fc and antibodies were purified by protein A affinity columns (Cytiva, Marlborough, MA, USA), as directed by the manufacturers. The recombinant protein preparations were dialyzed against phosphate-buffered saline (PBS) and stored in aliquots at −80°C. Recombinant protein purity was confirmed by SDS-PAGE. EH3 and EH8 Fab fragments were generated by overnight papain digestion at 37°C using immobilized papain agarose (ThermoFisher Scientific). Fab were separated from Fc and uncleaved IgG by passage over protein A resin followed by size-exclusion chromatography on a Superose 6 10/300 column before being used in SPR binding and X-ray crystallography. Also, for X-ray crystallography experiments, the expression plasmid encoding for codon-optimized SARS-CoV-2 RBD (329-538) with a C-terminal hexa-histidine tag was transfected into Expi293™ GnTI- (Thermo Fisher) cells (1×10^6^ cells/mL) with PEI-Max. One-week post-transfection, the clarified supernatant was purified on Ni-NTA column (Cytiva), followed by size-exclusion chromatography on HiLoad 26/600 Superdex 200 (Cytiva) equilibrated with PBS buffered saline pH 7.4.

### SARS-CoV-2 RBD ELISA (enzyme-linked immunosorbent assay)

The SARS-CoV-2 RBD ELISA assay used was previously described^23^. Briefly, recombinant SARS-CoV-2 RBD (2.5 μg/mL), or bovine serum albumin (BSA) (2.5 μg/mL) as a negative control, were prepared in PBS and were adsorbed to plates (MaxiSorp; Nunc) overnight at 4°C. Coated wells were subsequently blocked with blocking buffer (Tris-buffered saline [TBS] containing 0.1% Tween20 and 2% BSA) for 1 h at room temperature. Wells were then washed four times with washing buffer (TBS containing 0.1% Tween20). CR3022 mAb (50 ng/mL) or plasma from SARS-CoV-2-infected or uninfected donors (1/100; 1/250; 1/500; 1/1000; 1/2000; 1/4000) were prepared in a diluted solution of blocking buffer (0.1% BSA) and incubated in the coated wells for 90 min at room temperature. Plates were washed four times with washing buffer followed by incubation with HRP-conjugated anti-Human IgG secondary Abs (Invitrogen) (prepared in a diluted solution of blocking buffer [0.4% BSA]) for 1 h at room temperature, followed by four washes. HRP enzyme activity was determined after the addition of 40 μL of a 1:1 mix of Western Lightning oxidizing and luminol reagents (Perkin Elmer Life Sciences). Light emission was measured with a LB942 TriStar luminometer (Berthold Technologies). Signal obtained with BSA was subtracted for each antibody and was then normalized to the signal obtained with CR3022 mAb present in each plate.

### Pseudovirus neutralization assay

293T-ACE2 target cells were infected with single-round luciferase-expressing lentiviral particles^23^. Briefly, 293T cells were transfected by the calcium phosphate method with the pNL4.3 R-E- Luc plasmid (NIH AIDS Reagent Program) and a plasmid encoding for SARS-CoV-2 Spike WT at a ratio of 5:4. Two days post-transfection, cell supernatants were harvested and stored at –80°C until use. 293T-ACE2 target cells were seeded at a density of 1 × 10^4^ cells/well in 96-well luminometer-compatible tissue culture plates (PerkinElmer) 24 h before infection. Recombinant viruses in a final volume of 100 μL were incubated with the indicated plasma dilution (1/50; 1/250; 1/1250; 1/6250; 1/31250) or semi-log diluted antibody concentrations for 1 h at 37°C and were then added to the target cells followed by incubation for 48 h at 37°C. Cells were lysed by the addition of 30 μL of passive lysis buffer (Promega) followed by one freeze-thaw cycle. An LB942 TriStar luminometer (Berthold Technologies) was used to measure the luciferase activity of each well after the addition of 100 μL of luciferin buffer (15 mM MgSO4, 15 mM KH2PO4 [pH 7.8], 1 mM ATP, and 1 mM dithiothreitol) and 50 μL of 1 mM d-luciferin potassium salt (Prolume). The neutralization half-maximal inhibitory dilution (ID50) represents the plasma dilution to inhibit 50% of the infection of 293T-ACE2 cells by recombinant viruses bearing the SARS-CoV-2 S glycoproteins. The neutralization half-maximal inhibitory concentration (IC50) represents the antibody concentration to inhibit 50% of the infection of 293T-ACE2 cells by recombinant viruses bearing the SARS-CoV-2 S glycoproteins.

### Isolation of RBD-specific B cells

PBMCs from the pediatric patient of interest (Patient 12) were isolated from 5 mL of fresh blood through a ficoll-Paque™ gradient. To detect RBD-specific B cells, we conjugated recombinant RBD proteins with Alexa Fluor 488 or Alexa Fluor 647 (Thermo Fisher Scientific) according to the manufacturer’s protocol. Two million isolated PBMCs were labelled with the following antibodies for flow cytometry sorting: RBD-Alexa Fluor 647, RBD-Alexa Fluor 488, anti-hCD19- PE (clone HIB19, Biolegend), anti-hIgG-BV786 (clone G18-145, BD Biosciences), anti-hCD3- BV395 (clone SK7, BD Biosciences), anti-hCD14-PE-Cy7 (clone M5E2, BD Biosciences), anti-hIgD-BV650 (clone IA6-2, BD Biosciences) and 7-AAD nucleic acid dye (BD Biosciences) as a viability marker. Gating strategy is depicted in **Figure S1**. A total of nine RBD+ CD19^+^ IgG^+^ CD3^-^ CD14^-^ IgD^-^ 7AAD^-^ cells were sorted using a BD FACSAria^TM^ Fusion as single cells (1 cell per well) in a 96 well plate.

### Library generation

A library of cDNA was produced for each of the sorted single cells to identify the BCR sequence. Cells were sorted in 96 well plates in lysis buffer and reverse transcription and cDNA pre-amplification was performed as described^76^, performing 23 cycles of pre-amplification. cDNA quality was assessed on a BioAnalyzer (Agilent) using a high-sensitivity chip. V(D)J amplification was performed in two steps as described in the V(D)J library preparation kit (10X Genomics) according to the manufacturer’s instructions and adding to the first amplification the enrichment primer 1 (5’ CTACACGACGCTCTTCCGATCTAGCAGTGGTATCAACGCA 3’) and the enrichment primer 2 (5’ CTACACGACGCTCTTCCGATCTAGC 3’) to the second amplification step. These primers are designed to bind to the template switching oligonucleotides used in the cDNA generation and add to the amplicon the sequence of the Illumina read 1 primer. 5 ng of cDNA was used as input to the target amplification and 11 PCR cycles were performed for the first enrichment and 13 cycles for the second enrichment. After SPRI cleanup, the targeted amplification result was assessed on a BioAnalyzer. Following amplification of the V(D)J regions from cDNA, library construction was performed following the V(D)J reagents kit protocol (10X Genomics) for fragmentation, end repair, A-tailing, adaptor ligation and index PCR. The resulting libraries were sequenced on both an Illumina Nextera and an Oxford Nanopore instrument.

### BCR sequencing and assembly

Amplicons were submitted to Oxford Nanopore sequencing using the native barcoding and ligation sequencing protocol sample preparation kits (Oxford Nanopore Technologies) prior to loading onto the FLO-MIN106 flowcell. Raw sequencing data was base-called and demultiplexed with Guppy version 4.0.14 using configuration file dna_r9.4.1_450bps_hac.cfg. Demultiplexed reads were filtered and assembled into draft contigs using MAFFT, which were then polished using 4 successive rounds of Minimap2 and RACON using parameters “--secondary=no” and “ -m 8 -x - 6 -g -8”, respectively. The resulting contigs were then subjected to Medaka consensus correction with default parameters. A final round of consensus polishing was performed with Illumina short reads using Nextpolish with default parameters. IgBLAST was used to assess the sequence identity of the assembled BCRs, the CDR3 sequence and the V(D)J genes.

### Cloning of anti-RBD antibody from identified sequences & antibody production

From the nine different RBD-specific B cells isolated from the patient, 7 were successfully sequenced. Two sequences were identical, leading to a total of 6 different sequences. These assembled BCR sequences were synthesized into GeneBlocks (IDT DNA) and then cloned in the pTRIOZ-hIgG1 plasmid (InvivoGen) using the restriction enzymes SgrAI (NEB) and BsiWI-HF (NEB) for the variable domain of light chain, as well as AgeI-HF (NEB) and NheI-HF (NEB) for the variable domain of heavy chain. A sanger sequencing has been performed to ensure the quality of the cloned sequences. Using these vectors, anti-RBD antibodies (EH1, EH2, EH3, EH5 and EH8) were then produced, purified, and validated.

### Flow cytometry analysis of cell-surface staining

Using the standard calcium phosphate method, 10 µg of Spike expressor and 2.5 µg of a green fluorescent protein (GFP) expressor (pIRES2-eGFP; Clontech) was transfected into 2 × 10^6^ HEK 293T cells. At 48h post transfection, 293T cells were stained with anti-Spike monoclonal antibodies CV3-25, CV3-1, EH3 or EH8 (5 μg/mL) or using the ACE2-Fc chimeric protein (20 μg/mL) for 45 min at 37°C. Alternatively, CEM.NKr-Spike cells were incubated with increasing concentrations of CV3-1, EH3 or EH8 (0.0025 – 5 µg/mL). Alexa Fluor-647-conjugated goat anti-human IgG (H + L) Abs (Invitrogen) were used as secondary antibodies to stain cells for 30 min at room temperature. The percentage of transfected or transduced cells (GFP+ cells) was determined by gating the living cell population based on viability dye staining (Aqua Vivid, Invitrogen). Samples were acquired on an LSRII cytometer (BD Biosciences) and data analysis was performed using FlowJo v10.5.3 (Tree Star).

### Cell-to-cell fusion assay

To assess cell-to-cell fusion, 2 × 10^6^ 293T cells were co-transfected with plasmid expressing HIV- 1 Tat (1μg) and a plasmid expressing SARS-CoV-2 Spike (4 μg) using the calcium phosphate method. Two days after transfection, Spike-expressing 293T (effector cells) were detached with PBS-EDTA 1mM and incubated for 1 h with indicated amounts of CV3-1, EH3 or EH8 NAbs at 37°C and 5% CO2. Subsequently, effector cells (1 × 10^4^) were added to TZM-bl-ACE2 target cells that were seeded at a density of 1 × 10^4^ cells/well in 96-well luminometer-compatible tissue culture plates 24 h before the assay. Cells were co-incubated for 6 h at 37°C and 5% CO2, after which they were lysed by the addition of 40 μL of passive lysis buffer (Promega) and one freeze-thaw cycle. An LB942 TriStar luminometer (Berthold Technologies) was used to measure the luciferase activity of each well after the addition of 100 μL of luciferin buffer (15 mM MgSO4, 15 mM KH2PO4 [pH 7.8], 1 mM ATP, and 1 mM dithiothreitol) and 50 μL of 1 mM d-luciferin potassium salt (Prolume).

### Antibody dependent cellular cytotoxicity (ADCC) assay

For evaluation of anti-SARS-CoV-2 ADCC activity, parental CEM.NKr CCR5+ cells were mixed at a 1:1 ratio with CEM.NKr-Spike cells. These cells were stained for viability (AquaVivid; Thermo Fisher Scientific) and a cellular dye (cell proliferation dye eFluor670; Thermo Fisher Scientific) and subsequently used as target cells. Overnight rested PBMCs were stained with another cellular marker (cell proliferation dye eFluor450; Thermo Fisher Scientific) and used as effector cells. Stained effector and target cells were mixed at a 10:1 ratio in 96-well V-bottom plates. Titrated concentrations of CV3-1, EH3 or EH8 mAbs were added to the appropriate wells. The plates were subsequently centrifuged for 1 min at 300xg, and incubated at 37°C, 5% CO2 for 5 h before being fixed in a 2% PBS-formaldehyde solution. ADCC activity was calculated using the formula: [(% of GFP+ cells in Targets plus Effectors) - (% of GFP+ cells in Targets plus Effectors plus antibody)]/(% of GFP+ cells in Targets) × 100 by gating on transduced live target cells. All samples were acquired on an LSRII cytometer (BD Biosciences) and data analysis performed using FlowJo v10.5.3 (Tree Star).

### X-ray crystallography of EH3 and EH8 with SARS-CoV-2 RBD

Fabs of EH3 and EH8 were prepared by standard papain digestion as described^25^. The purified Fabs were mixed with excessive SARS-CoV-2 RBD (319-537, molar ratio ∼1:4) respectively and incubated at 4 °C overnight before separation on HiLoad 26/600 Superdex 200 column (Cytiva) which was pre-equilibrated in 10mM Tris pH 8.0 and 100mM ammonium acetate. The complex fractions were pooled and concentrated to >10 mg/mL. Crystallization trials of the Fab-RBD complexes were performed by mixing 1:1 ratio of protein to well solution in the vapor-diffusion hanging drop manner. After 2 weeks incubation at 21 °C, diffraction-quality crystals of EH3-RBD and EH8-RBD were obtained in 0.1 M sodium citrate pH 5.0, 8% w/v PEG 8000 and 0.2 M sodium chloride, 0.1 M HEPES pH 7.5, 12% w/v PEG 8000, respectively. Crystals were cryo-cooled by liquid nitrogen in the crystallization condition supplemented with 20% 2-methyl-2, 4-pentanediol (MPD) used as the cryoprotectant. X-ray diffraction data was collected at the SSRL beamline 9-2 and was indexed, integrated, scaled with HKL3000 ^77^. The structure was solved by molecular replacement in Phenix using 7S4S (template for RBD) and 7NAB (for Fabs) as independent searching models. Iterative cycles of model building and refinement were done in Coot ^78^ and Phenix ^79^. Data collection and refinement statistics are provided in **Table S2**.

### smFRET imaging of S on Lentivirus VLPs

smFRET imaging of SARS-CoV-2 SWT expressed on the surface of HIV-1 VLPs and prepared as previously described ^24^. Two short peptides labeling tags (Q3: GQQQLG; A4: DSLDMLEM) were introduced into designed positions in the S1 subunit on the plasmid encoding SWT. With ∼1:20 plasmid ratio of tagged-S and unlabeled SWT, more than 95% S trimers present on VLP surface had one dual-tagged protomer and two unlabeled protomers. Viral particles were harvested 40 h post-transfection, filtered with a 0.45 mm pore size filter, and purified using ultra-centrifugation at 25,000 rpm for 2 h through a 15% sucrose cushion made in PBS. The resulting viral particles were re-suspended in 50 mM pH 7.5 HEPES buffer, labeled with self-healing Cy3 and Cy5 derivatives (LD555-CD and LD655-CoA, respectively) and purified through an Optiprep™ (Sigma Aldrich) gradient as previously described ^24^. smFRET images of viral particles was acquired on a home-built prism-based total internal reflection fluorescence (TIRF) microscope. The conformational effects of 200 µg/mL soluble ACE2, EH3 or EH8 IgGs on SARS-CoV-2 Spike were tested by pre-incubating fluorescently labeled viruses for 60 min at room temperature before imaging in the continued presence of the antibodies. Signals were simultaneously recorded on two synchronized ORCA-Flash4.0 V3 sCMOS cameras (Hamamatsu) at 25 frames per second for 80 s smFRET data analysis was performed using MATLAB (MathWorks)-based customized SPARTAN software package. Each FRET histogram was fitted into the sum of four Gaussian distributions in Matlab, where each Gaussian distribution represents one conformation and the area under each Gaussian curve estimates the occupancy of each state.

### Surface plasmon resonance

All surface plasma resonance assays were performed on a Biacore 3000 (Cytiva) with a running buffer of 10 mM HEPES pH 7.5 and 150 mM NaCl supplemented with 0.05% Tween 20 at 25 °C. For the kinetic measurements of SARS-CoV-2 VOCs’ RBDs to EH3 and EH8, either of the IgGs (∼250-350 response unit, RU) was first immobilized on a protein A chip (Cytiva) and 2-fold serial dilutions of the RBDs were then injected with concentrations ranging from 6.25 to 200 nM. After each cycle the sensor chip was regenerated with 0.1 M Glycine pH 2.5. All sensorgrams were corrected by subtraction of the corresponding blank channel in addition to the buffer background and the kinetic constant determined using a 1:1 Langmuir model with the BIAevaluation software (Cytiva). Goodness of fit of the curve was evaluated by the Chi^2^ value with a value below 3 considered acceptable. The sensorgrams are shown in **Figures S2A-B** and the kinetic constants are summarized in **Table 1**.

### Biolayer interferometry

Binding kinetics were performed on using an Octet RED96e system (ForteBio) at 25 °C with shaking at 1,000 RPM. Amine Reactive Second Generation (AR2G) biosensors (Sartorius) were hydrated in water, then activated for 300 s with a solution of 5 mM sulfo-NHS and 10 mM EDC prior to amine coupling. Either SARS-CoV-2 RBD WT or the E484K mutant were loaded into AR2G biosensor at 12.5 µg/mL at 25°C in 10 mM acetate solution pH 5 for 600 s then quenched into 1 M ethanolamine solution pH 8.5 for 300 s. Loaded biosensor were placed in 10X kinetics buffer for 120 s for baseline equilibration. Association of EH3 or EH8 IgG (in 10X kinetics buffer) to the different RBD proteins was carried out for 180 s at various concentrations in a two-fold dilution series from 50 nM to 3.125 nM prior to dissociation for 300 s. The data were baseline subtracted prior to fitting performed using a 1:1 binding model and the ForteBio data analysis software. Calculation of on rates (kon), off rates (koff), and affinity constants (KD) was computed using a global fit applied to all data.

### Radioactive labeling and immunoprecipitation

For pulse-labeling experiments, 5 × 10^5^ 293T cells were transfected by the calcium phosphate method with SARS-CoV-2 Spike expressors. One day after transfection, cells were metabolically labeled for 16 h with 100 μCi/mL [^35^S] methionine-cysteine ([^35^S] protein labeling mix; PerkinElmer) in Dulbecco’s modified Eagle’s medium lacking methionine and cysteine and supplemented with 10% of dialyzed fetal bovine serum and 1X GlutaMAX™ (ThermoFisher Scientific). Simultaneously, cells were treated with or without ACE2-Fc (20 µg/mL) CV3-1, EH3, EH8, CV3-25, Casirivimab or Imdevimab (5 µg/mL). Cells were subsequently lysed in radioimmunoprecipitation assay (RIPA) buffer (140 mM NaCl, 8 mM Na2HPO4, 2 mM NaH2PO4, 1% NP-40, 0.05% sodium dodecyl sulfate [SDS], 1.2 mM sodium deoxycholate [DOC]) with protease inhibitors (ThermoFisher Scientific). Precipitation of radiolabeled SARS-CoV-2 Spike glycoproteins from cell lysates or supernatant was performed with CV3-25 in combination with a polyclonal rabbit antiserum raised against SARS-CoV-2 RBD protein for 1 h at 4°C in the presence of 45 μL of 10% protein A-Sepharose beads (Cytiva).

### *In vivo* prophylactic efficacy of EH3 in Delta-nLuc VOC challenged K18-hACE2 mice

For in vivo efficacy analyses, EH3 IgG were intraperitoneally (*i.p.*) administered at 12.5 mg/kg to K18-hACE2 mice under prophylaxis regimen. 12 h post-administration, the mice were challenged intranasally (*i.n.*) with 1 × 10^5^ PFU SARS-CoV-2 Delta-nLuc VOC. in 25-30 µL volume under anesthesia (0.5 - 5 % isoflurane) delivered using precision Dräger vaporizer with oxygen flow rate of 1 L/min). The starting body weight was set to 100 %. For survival experiments, mice were monitored every 6-12 h starting six days after virus administration. Lethargic and moribund mouse or mouse that had lost more than 20 % of their body weight, were sacrificed and considered to have succumbed to infection for Kaplan-Meier survival plots.

### Bioluminescence Imaging (BLI) of SARS-CoV-2 infection

All standard operating procedures and protocols for IVIS imaging of SARS-CoV-2 infected animals under ABSL-3 conditions were approved by IACUC, IBSCYU and YARC. All the imaging was carried out using IVIS Spectrum® (PerkinElmer) in XIC-3 animal isolation chamber (PerkinElmer) that provided biological isolation of anesthetized mice or individual organs during the imaging procedure. All mice were anesthetized via isoflurane inhalation (3 - 5% isoflurane, oxygen flow rate of 1.5 L/min) prior and during BLI using the XGI-8 Gas Anesthesia System. Prior to imaging, 100 μL of Nanoluc luciferase substrate, furimazine (NanoGlo™, Promega, Madison, WI) diluted 1:40 in endotoxin-free PBS was retro-orbitally administered to mice under anesthesia. The mice were then placed into XIC-3 animal isolation chamber (PerkinElmer) pre- saturated with isothesia and oxygen mix. The mice were imaged in both dorsal and ventral position at indicated days post infection. Whole-body was imaged again after euthanasia and necropsy by spreading additional 200 μL of substrate on to exposed intact organs. Identified infected areas of interest were isolated, washed in PBS to remove residual blood and placed onto a clear plastic plate. Additional droplets of furimazine in PBS (1:40) were added to isolated organs and soaked in substrate for 1-2 min before BLI.

Images were acquired and analyzed using Living Image v4.7.3 *in vivo* software package (Perkin Elmer Inc). Image acquisition exposures were set to auto, with imaging parameter preferences set in order of exposure time, binning, and f/stop, respectively. Images were acquired with luminescent f/stop of 2, photographic f/stop of 8. Binning was set to medium. Comparative images were compiled and batch-processed using the image browser with collective luminescent scales. Photon flux was measured as luminescent radiance (p/sec/cm2/sr). During luminescent threshold selection for image display, luminescent signals were regarded as background when minimum threshold levels resulted in displayed radiance above non-tissue-containing or known uninfected regions.

### Measurement of viral burden

Indicated organs (nasal cavity, brain and lungs) from infected or uninfected mice were collected, weighed, and homogenized in 1 mL of serum free RPMI media containing penicillin-streptomycin and homogenized in 2 mL tube containing 1.5 mm Zirconium beads with BeadBug 6 homogenizer (Benchmark Scientific, TEquipment Inc). Virus titers were measured using two highly correlative methods. First, the total RNA was extracted from homogenized tissues using RNeasy plus Mini kit (QIAGEN), reverse transcribed with iScript advanced cDNA kit (Bio-Rad) followed by a SYBR Green Real-time PCR assay for determining copies of SARS-CoV-2 N gene RNA using primers SARS-CoV-2 N F: 5’-ATGCTGCAATCGTGCTACAA-3’ and SARS-CoV-2 N R: 5’-GACTGCCGCCTCTGCTC-3’. RNA extracted from the corresponding tissues of uninfected mice were used for normalization of N mRNA copy numbers. We observe a Ct value ranging between 38 and 40 in uninfected samples with our primer set and conditions. Therefore, we used a Ct value of 38 (lower end) to normalize all the data for estimating N RNA levels.

Second, serially diluted clarified tissue homogenates were used to infect Vero-E6 cell culture monolayer. We used Nanoluc luciferase activity as a shorter surrogate for plaque assay. Infected cells were washed with PBS and then lysed using 1X Passive lysis buffer. The lysates transferred into a 96-well solid white plate (Costar Inc) and Nanoluc luciferase activity was measured using Tristar multiwell Luminometer (Berthold Technology, Bad Wildbad, Germany) for 2.5 s by adding 20 μL of Nano-Glo® substrate in assay buffer (Promega Inc, WI, USA). Uninfected monolayer of Vero cells treated identically served as controls to determine basal luciferase activity for obtaining normalized relative light units. The data were processed and plotted using GraphPad Prism v8.4.3.

### Analyses of signature inflammatory cytokines mRNA expression

Brain, lung and nose samples were collected from mice at the time of necropsy. Approximately, 20 mg of tissue was suspended in 500 μL of RLT lysis buffer, and RNA was extracted using RNeasy plus Mini kit (Qiagen), reverse transcribed with iScript advanced cDNA kit (Bio-Rad). To determine levels of signature inflammatory cytokines, multiplex qPCR was conducted using iQ Multiplex Powermix (Bio Rad) and PrimePCR Probe Assay mouse primers FAM-GAPDH, HEX-IL6, TEX615-CCL2 and Cy5-CXCL10. The reaction plate was analyzed using CFX96 touch real time PCR detection system. Scan mode was set to all channels. The PCR conditions were 95°C 2 min, 40 cycles of 95°C for 10 s and 60°C for 45 s, followed by a melting curve analysis to ensure that each primer pair resulted in amplification of a single PCR product. mRNA levels of Il6, Ccl2 and Cxcl10 in the cDNA samples of infected mice were normalized to Gapdh with the formula ΔCt(target gene) = Ct(target gene)-Ct(gapdh). The fold increase was determined using 2−ΔΔCt method comparing treated mice to uninfected controls.

### Quantification and Statistical Analysis

Data were analyzed and plotted using GraphPad Prism software (San Diego, CA). *P* values lower than 0.05 were considered statistically significant. *P* values were indicated as ∗, *P* < 0.05; ∗∗, *P* < 0.01; ∗∗∗, *P* < 0.001; ∗∗∗∗, *P* < 0.0001.

## Supplemental information

**Table S1.** Summary of biolayer interferometry kinetic constants. Related to Table 1 and Figure S2C.

| Analyte | Immobilized ligand | Fitting mode | $k_a$ (M<sup>-1</sup>s<sup>-1</sup>) | $k_d$ (s<sup>-1</sup>) | K<sub>D</sub> (pM) | K<sub>D</sub> reduction fold compared to RBD<sub>WT</sub> |
| --- | --- | --- | --- | --- | --- | --- |
| EH3 IgG | RBD<sub>WT</sub> | 1:1 | 1.01 x 10<sup>5</sup> | 6.09 x 10<sup>-7</sup> | 6.06 | 1 |
|  | RBD<sub>E484K</sub> | 1:1 | 1.11 x 10<sup>5</sup> | 4.46 x 10<sup>-7</sup> | 4.04 | 0.67 |
| EH8 IgG | RBD<sub>WT</sub> | 1:1 | 2.11 x 10<sup>5</sup> | 3.55 x 10<sup>-7</sup> | 1.68 | 1 |
|  | RBD<sub>E484K</sub> | 1:1 | 1.83 x 10<sup>5</sup> | 6.01 x 10<sup>-7</sup> | 3.29 | 1.96 |

**Table S2.**
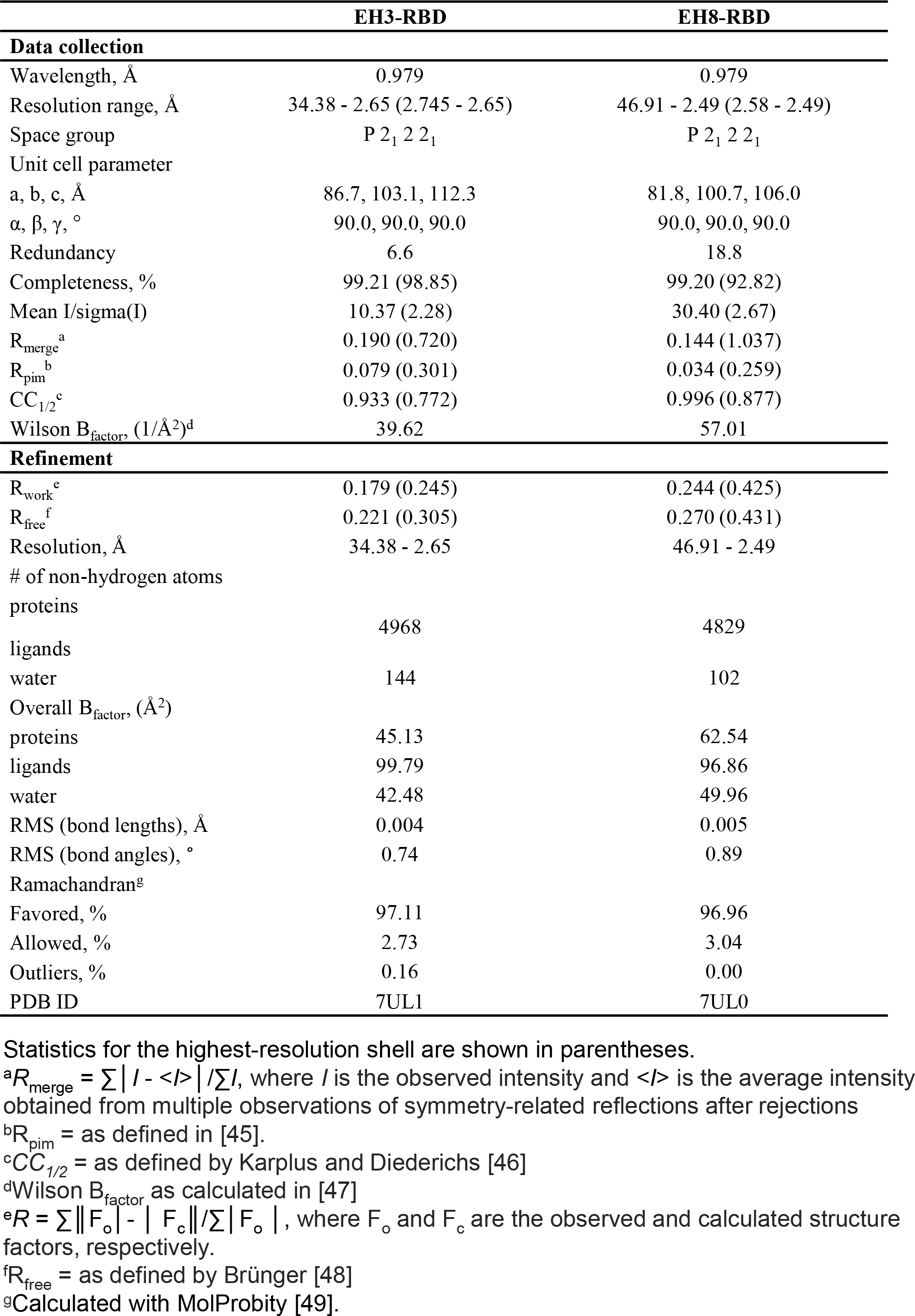
Data collection and model refinement statistics. Related to Figure 3.

**Figure S1.**
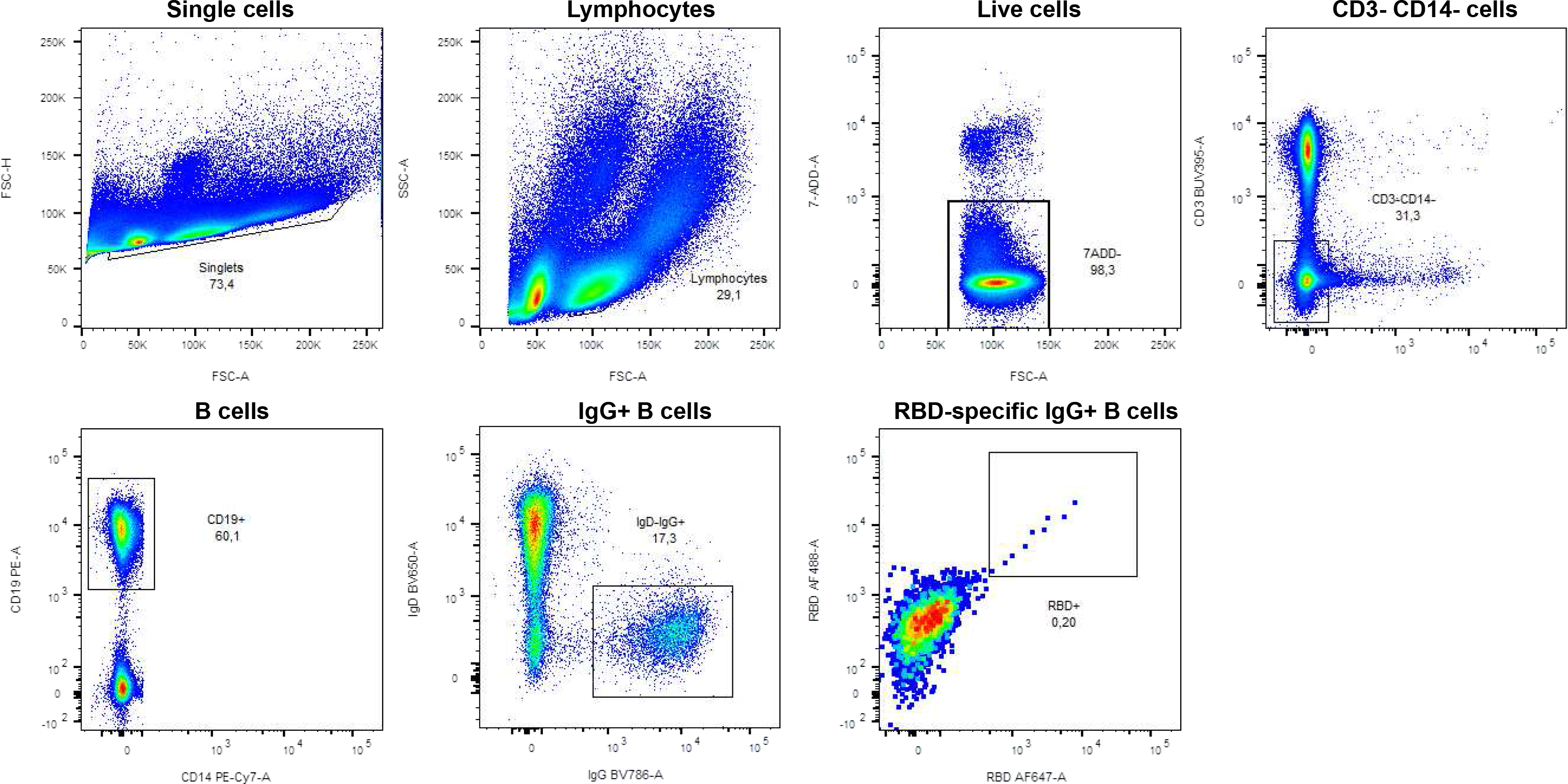
Gating strategy for the isolation of RBD-specific mAbs. Related to Figure 1. Flow cytometry gates to identify RBD-specific B cells from PBMCs of a pediatric patient (Patient 12). RBD-specific B cells were identified according to cell morphology by light-scatter parameters and excluding doublets cells while gating on live cells (7-AAD-). Cells were then gated based on lineage markers: CD3- (T cell marker), CD14- (monocyte marker) and CD19+ (B cell marker). Cells were further gated for isotypic expression: IgD- (naïve) and IgG+ (class-switched). Finally, RBD-specific IgG+ B cells were identified using a dual staining with fluorescent RBD probes (RBD-AF647 and RBD-AF488). The nine RBD-specific B cells were sorted as single cells to clone their BCR sequence.

**Figure S2.**
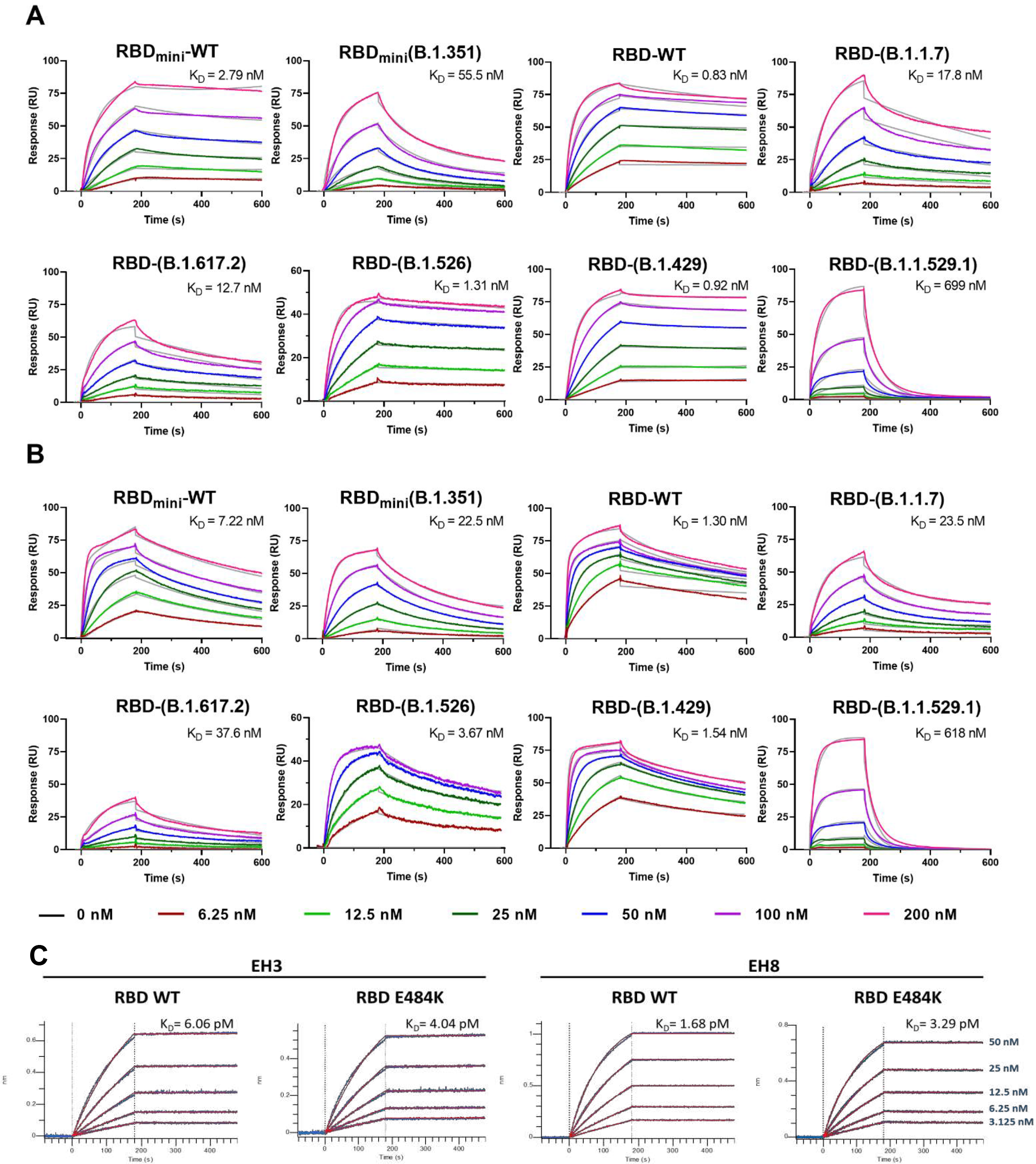
Binding affinity of RBDs of SARS-CoV-2 VOCs to EH3 and EH8. Related to Table 1. (**A-B**) SPR kinetic measurement of SARS-CoV-2 RBDs binding to immobilized (**A**) EH3 IgG and (**B**) EH8 IgG. Either IgGs (∼250-350 RU) were immobilized on a Protein A chip and 2-fold dilutions of SARS-CoV-2 RBDs of WT and 6 VOCs (6.25-200nM) were injected as flow analytes. The experimental sensorgrams are colored in indicated colors and the 1:1 Langmuir fitting are in grey. RBD_mini_ is the segment of residue 329-527 and otherwise are residue 319-537. The detailed SPR kinetic constants are listed in **Table 1**. (**C**) Binding kinetics between SARS-CoV-2 RBD (WT or E484K) and EH3 or EH8 mAbs assessed by biolayer interferometry. Biosensors loaded with RBD proteins were soaked in two-fold dilution series of indicated mAbs (50 nM**–**3.125 nM) at 25°C. Raw data are shown in blue and 1:1 binding model is shown in red. The detailed biolayer interferometry kinetic constants are listed in **Table S1**.

**Figure S3.**
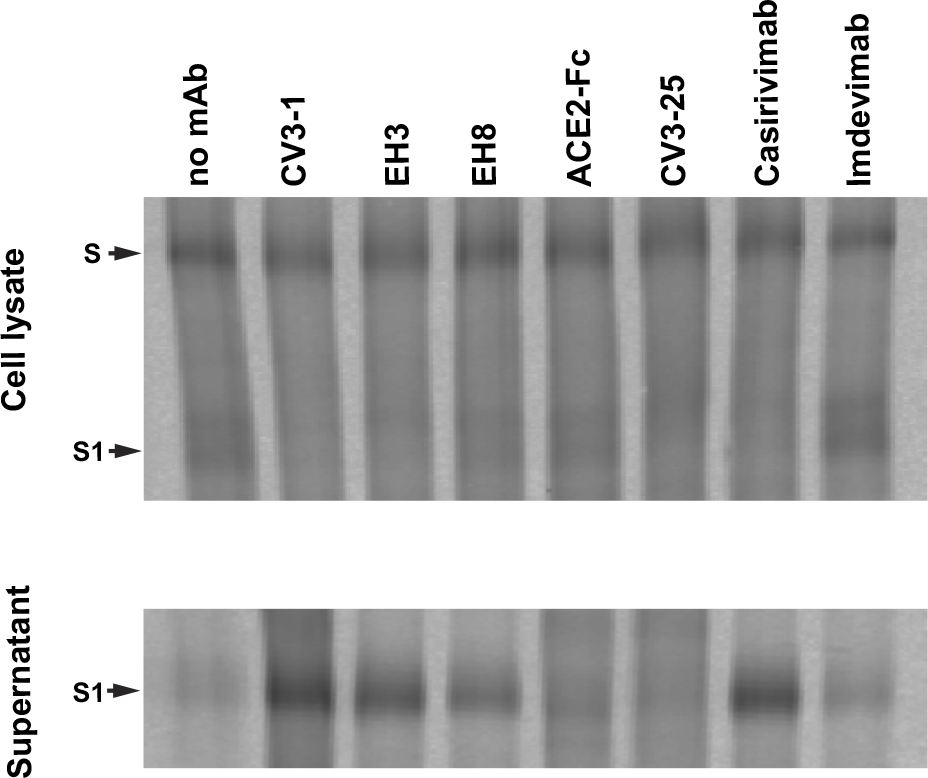
The ability of RBD-specific mAbs to induce S1 shedding is epitope-dependent. Related to Figure 4. S1 shedding was evaluated by transfection of 293T cells to express SARS-CoV-2 S D614G followed by radiolabeling using 35S-methionine/cysteine mix in presence of RBD-specific mAbs (CV3-1, EH3, EH8), ACE2-Fc, S2-specific CV3-25, or two FDA-approved RBD-specific mAbs (Casirivimab and Imdevimab). This was followed by immunoprecipitation of cell lysates and supernatant with CV3-25 and a rabbit antiserum raised against SARS-CoV-2 RBD produced in-house. Samples were analyzed by SDS-PAGE and autoradiography.

**Figure S4.**
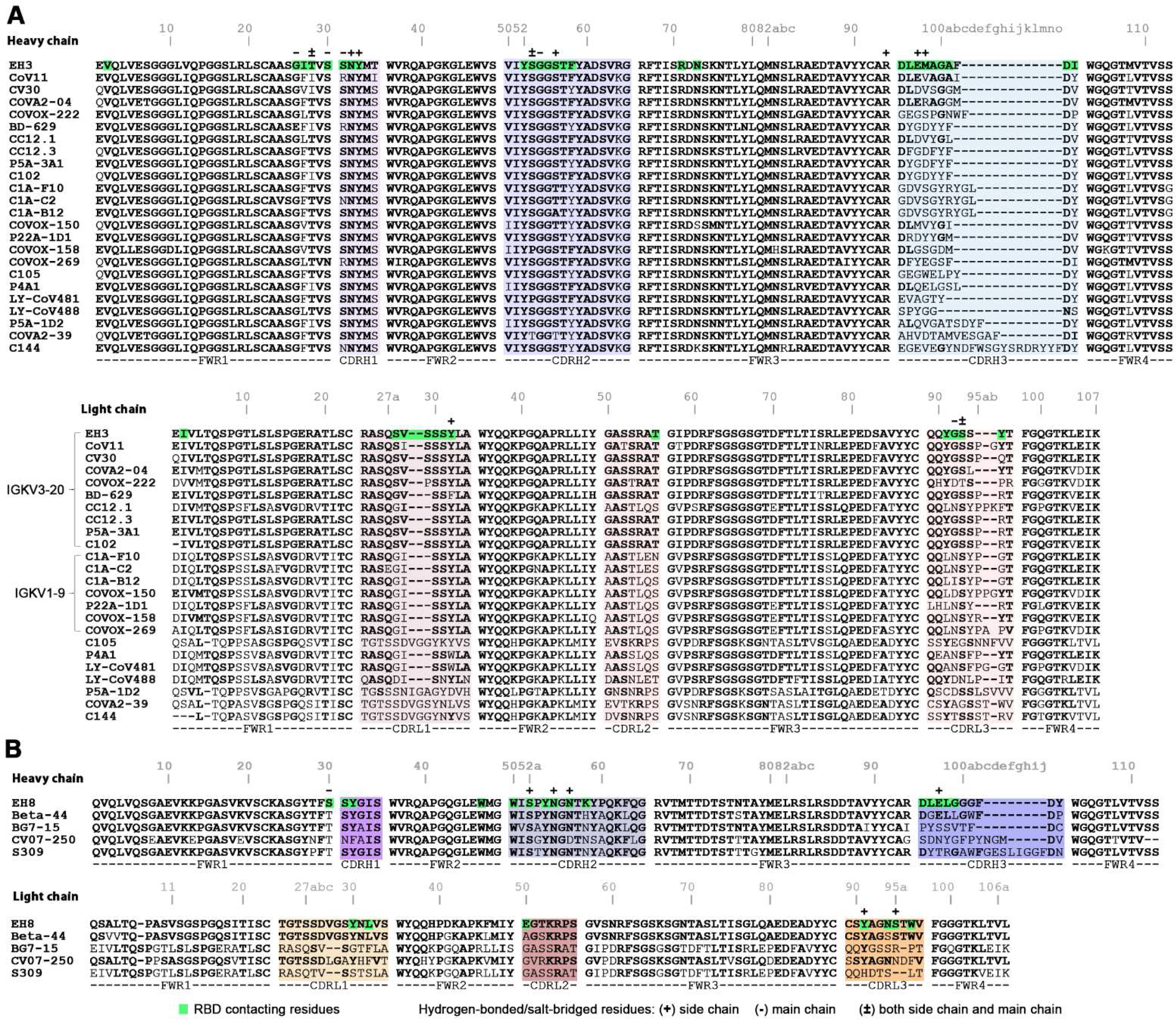
V_H_ and V_L_ sequence alignments of EH3 and EH8 to other structurally available RBD-specific mAbs. Related to Figure 4. (**A**) Alignments of EH3 and other 23 IGHV3-53 encoded Abs. (**B**) Alignments of EH8 and other 4 IGHV1-18 Abs. The RBD-contacting residues (BSA>0) in EH3-RBD and EH8-RBD crystal structures are shaded in green. Residues involved in salt-bridges or H-bonds to the RBD are marked above the sequence with (+) for the side chain, (-) for the main chain and (±) for both.

